# Continuous, high-speed 3D topographic profiling of living tissue monolayers

**DOI:** 10.64898/2026.07.23.740215

**Authors:** Daeeun Shin, Jimin Park, Jihye Ahn, Andrew Choi, Joon Ho Kang

## Abstract

Tissue monolayers are active living matter whose three-dimensional (3D) architecture and mechanical remodeling regulate tissue function, yet fast and continuous topographic monitoring remains difficult. Here, we present FLuorescence EXclusion microscopy fOr Monolayers (FLEXOM), a microfluidic platform that combines multilayer micropillar arrays with negative-staining optics to convert wide-field images into self-calibrated 3D height maps. Geometry-anchored reference cavities maintain in-frame calibration standards even in fully confluent monolayer fields, enabling continuous 3D profiling from isolated cells to confluent sheets with sub-second temporal resolution (~250 ms), sub-micrometer axial precision (~0.37 µm), and multi-day biocompatibility and operational stability (>96 h). FLEXOM reveals two distinct anisotropic signatures in tissue monolayers. First, monolayers of different cell types show distinct vertical thickness profiles despite comparable lateral footprints. Only two vertical descriptors, mean height and height variability, distinguished cell types more accurately than nine 2D lateral descriptors, including area, circularity, and convexity (95% vs. 76%). Second, under acute osmotic shock, confluent monolayers exhibit geometrically anisotropic and temporally decoupled vertical–lateral remodeling: height changes dominate the overall volume response and precede lateral remodeling. Surprisingly, cyclic isotonic–hypotonic pulses every 3 min confine remodeling entirely to the vertical axis. Overall, our work provides a high-resolution 3D profiling tool and a framework for axis-resolved analysis of topographic responses in active living matter.

## 1. Introduction

Tissue monolayers, such as epithelial and endothelial sheets, behave like active soft matter, displaying self-assembly and continuous remodeling of their own architecture [1,2]. Unlike passive materials, living matter actively consumes metabolic energy and reshapes itself in response to internal signals and external cues, intrinsically coupling their morphology and mechanics [3–8]. Consequently, even subtle topographic variations in three-dimensional architecture can redistribute local stresses and drive collective behaviors, including directed tissue migration [9,10], cell-fate decisions [5,11,12], differentiation [12,13], homeostatic remodeling [14–16], and wound healing [17]. Tissue topography is therefore not merely a morphological signature, but a dynamic material-state variable that emerges from the coupling between mechanics and function. Understanding how tissue monolayer mechanics and function are coupled requires real-time, quantitative measurement of their full 3D topography.

Yet, analysis of tissue monolayer architecture remains largely confined to two dimensions (2D). Two-dimensional descriptors derived from planar (xy) imaging [18,19], such as projected area [12] or aspect ratio [10], cannot resolve dynamics along the vertical (z) axis, including apical height fluctuations and extrusion or intrusion events [20,21]. Several modalities add axial resolution to 2D measurements and enable 3D topographical reconstruction [22–24], but continuous long-term monitoring of living matter remains a major challenge [25]. Confocal and light-sheet microscopy typically require fluorescent labeling that imposes phototoxicity and z-stack acquisition incompatible with long-term observation [25,26]. Atomic force microscopy is label-free but mechanically invasive on soft biological samples; the scanning probe deforms the specimen and throughput remains low [27]. Holographic [28] and quantitative phase imaging [29–31] offer label-free, non-invasive 3D reconstruction, yet most implementations have been limited to isolated or suspended cells and require heavy computational resources. Fluorescence exclusion microscopy (FXm) [32] offers an alternative route that simultaneously satisfies the criteria for high-speed, label-free, and non-invasive imaging. In FXm, the sample is not directly labeled; instead, it excludes fluorescent dye from the surrounding medium, producing a negative-contrast signal that can be converted into local axial thickness from a single wide-field image. Yet, FXm has remained largely limited to isolated single cells [32–35], due to the following three technical barriers that must be addressed simultaneously for extending its application to confluent tissues: (i) the lack of long-term stable *in situ* calibration references to convert fluorescence intensity into absolute axial height (µm); (ii) the absence of rapid, continuous fluid exchange within the chamber to suppress photobleaching-driven drift and maintain culture medium during long-term imaging; and (iii) the lack of a robust single-cell segmentation scheme for densely packed sheets.

Here, we present FLEXOM (FLuorescence EXclusion microscopy fOr Monolayers), an architected microfluidic platform designed for label-free, spatiotemporal profiling of living tissue monolayer topography. FLEXOM enables full 3D topographic dynamics to be continuously tracked at the single-cell level at frame intervals as short as 250 ms and over more than 4 days (>96 h). Using this approach, we resolved cell-type dependent topographic signatures at confluence and discovered an emergent, minute-scale anisotropic volume regulation in monolayers that cannot be captured by conventional 2D imaging. By making fast, quantitative, and long-term 3D topographic profiling accessible in a simple wide-field format, FLEXOM enables high-throughput mapping of structure–function relationships in engineered bio-interfaces and active living materials.

## 2. Results

### 2.1 In situ self-calibration via multilayered micropillar arrays

The central challenge in negative-contrast imaging (or FXm) of confluent tissues is the loss of a stable zero-height reference once cells tile the entire field of view. FLEXOM overcomes this challenge by embedding multilayered micropillar arrays into the microfluidic channel to achieve *in situ*, frame-by-frame self-calibration for fluorescence-to-height conversion (Figure 1a, Figure S1a). At each pixel (x, y), the local cell height *h*_*x,y*_ is calculated from a linear intensity-to-height relationship anchored by two geometric reference intensities (Figure 1b).

**FIGURE 1.**
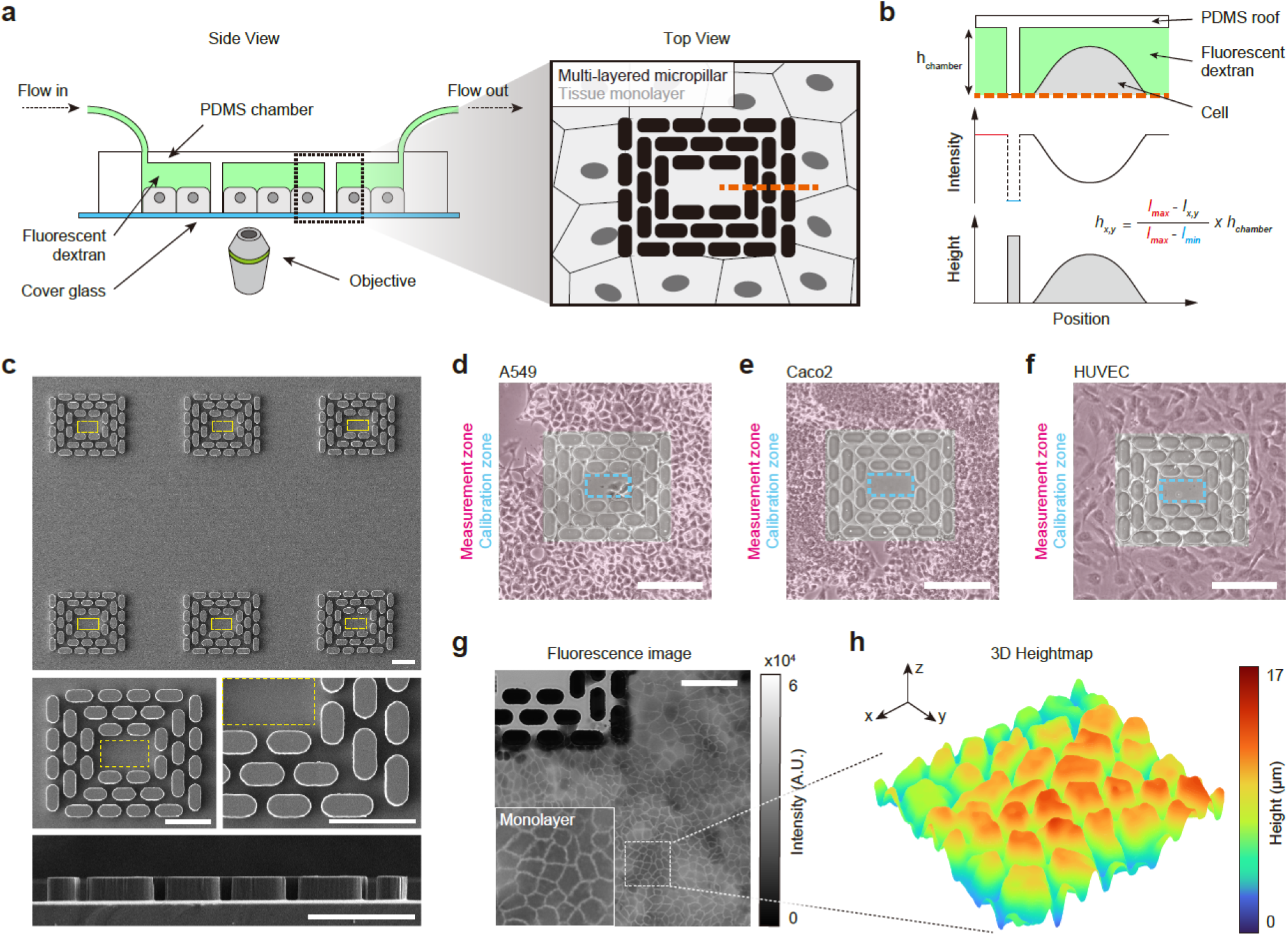
FLEXOM: Label-free 3D topographic profiling of living tissue monolayers via *in situ* self-calibrated fluorescence exclusion. (a) Schematic of the FLEXOM platform. Cells are cultured in a PDMS microfluidic chamber integrated with a multilayered micropillar array, under continuous perfusion of fluorescent dextran-supplemented medium (Side view, left; Top view, right). (b) Working principle of the fluorescence exclusion-based height calibration along the selected cross section (orange dashed line in a). Pixel-wise fluorescence intensity is topographically converted to absolute height using two internal reference signals extracted frame-by-frame: *I*_*max*_ (red) derived from the cell-free void zone and *I*_*min*_ (blue) from the PDMS pillar. (c) Scanning electron microscopy (SEM) images of the multilayer micropillar structure, providing geometry-anchored, cell-free reference zones (yellow dashed boxes) for *in situ* self-calibration. Scale bars, 100 µm. (d-f) Representative phase images of A549 (d), Caco-2 (e), and HUVEC (f) tissue monolayers after 4 days of culture within the platform. Red shading, measurement zone; blue dashed line, calibration zone. Scale bars, 200 μm. (g) Representative wide-field fluorescence image of a confluent A549 tissue monolayer. Scale bar, 200 µm. (h) 3D height map reconstructed from the selected region (white dashed box in g) by per-pixel application of the calibration in (b).

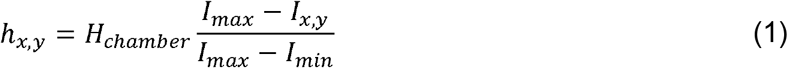

where *I*_*max*_ is the maximum intensity measured in the cell-free void zone (red dashed line in Figure 1b), representing a fully dextran-filled chamber with no cell in the optical path. This condition corresponds to a cell height of 0□µm. *I*_*min*_ is the minimum intensity measured above a full-height PDMS pillar (blue dashed line in Figure 1b), corresponding to complete dextran exclusion over the chamber height (*h*_*chamber*_ ≈ 27 µm). Frame-by-frame extraction of *I*_*max*_ and *I*_*min*_ cancels out slow illumination drift and minor frame-to-frame variations in camera acquisition time, enabling robust and precise 3D topographic reconstruction.

The multilayered micropillar array maintains these two reference regions within the imaging field during confluent tissue growth. The full-height PDMS pillars provide the dextran-excluded regions used to define *I*_*min*_. An enclosed cell-free cavity within the same array provides the dextran-filled region used to define *I*_*max*_ (Figure 1c). The cavity is bordered by densely spaced 5 µm-pitch pillars (yellow dashed boxes in Figure 1c), which restrict cell entry while allowing fluorescent dextran exchange with the surrounding medium. As a result, both calibration anchors remain accessible throughout long-term imaging, independent of tissue confluence in the surrounding measurement region. The array therefore functions as an integrated calibration scaffold for quantitative, frame-by-frame height reconstruction in confluent monolayers.

We also interfaced the microfluidic chip with a custom pneumatic controller to maintain stable physiological conditions during long-term culture and live imaging (Figure S1b). The controller regulates chamber pressure and perfusion flow rate in a closed-loop manner, continuously delivering fresh culture medium equilibrated at 37 °C and 5% CO_2_. Programmed, intermittent fluidic control minimizes photobleaching by periodically flushing the chamber with fresh, unbleached medium, while maintaining minimal shear stress under low-pressure perfusion conditions (ΔP ≤ 1 kPa; Figure S2).

Biocompatibility and stability assays confirmed that FLEXOM remains fully operational over 96 h of continuous imaging in three representative tissue-forming cell types (A549, human lung epithelial carcinoma; Caco-2, human intestinal epithelial; HUVEC, primary human umbilical vein endothelial), with no detectable impact on cell viability or calibration integrity across the full course of tissue monolayer assembly (Figure 1d–f, Figure S3). Notably, a high-molecular-weight fluorescent dye (2,000 kDa FITC-dextran) was not endocytosed and did not alter cell growth or morphology, supporting its suitability as a stable contrast agent (Figure S4). With validated calibration stability and tissue compatibility, FLEXOM yields long-term stable, quantitative 3D topographic maps directly from wide-field fluorescence (Figure 1g,h), supporting its use as a label-free 3D topographic profiler compatible with diverse tissue monolayers.

### 2.2 Accuracy, precision, and stability of FLEXOM measurements

We next asked how accurate, precise, and stable the FLEXOM 3D topographic measurements are (Figure 2). We first confirmed that FLEXOM resolves a known microscale topography by imaging the embedded multilayered micropillar array (Figure 2a), whose geometry was independently defined by CAD design and SEM imaging (Figure 1c, Figure S1a). FLEXOM reconstructed the array as a discrete 3D height structure with well-resolved pillar levels (Figure 2b), and cross-sectional profiles further resolved the pillar heights and spacing expected from the designed multilayered geometry (Figure 2c).

**FIGURE 2.**
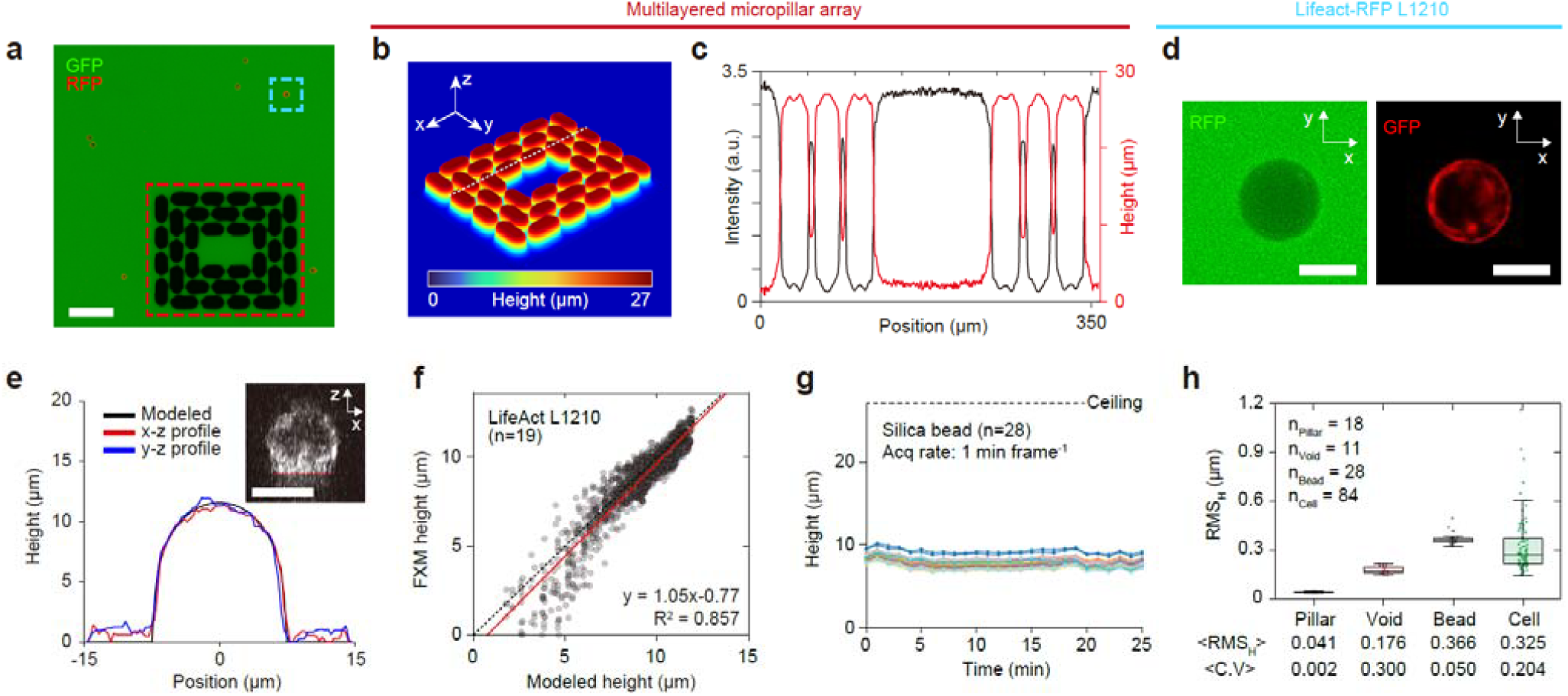
Quantitative validation and sub-micrometer axial precision of FLEXOM height measurements. (a) Representative overlaid fluorescence image showing FLEXOM signal (GFP), LifeAct-RFP L1210 cells (RFP; blue dashed box), and a multilayered micropillar array (red dashed box). Scale bar, 100 μm. (b) 3D reconstruction of the multilayered micropillar array (red dashed box in a). Color encodes absolute height. (c) Cross-sectional profiles of fluorescence intensity (black) and reconstructed height (red) along the white dashed line in (b). (d) Magnified fluorescence images of the representative LifeAct-RFP L1210 cell (blue dashed box in a), showing FLEXOM signal (GFP, left) and LifeAct-RFP signal (RFP, right). Scale bars, 10 μm. (e) Cross-sectional height profiles of a representative LifeAct-RFP L1210 cell measured along the x–z (red) and y–z (blue) directions, compared with the modeled profile (black). Inset, representative confocal x–z section showing basal adhesion of a LifeAct-RFP L1210 cell to the substrate (red arrow). Scale bar, 10 μm. (f) Correlation between FLEXOM-measured and modeled heights for LifeAct-RFP L1210 cells (n = 19 cells). Red line, linear regression (y = 1.05x −0.77); black dashed line, identity (y = x). Each dot represents a matched pair at a single position along the cross-section al profile. (g) Temporal height traces of individual silica beads (n = 28 beads; each line represents one bead). (h) RMS_H_ distributions for pillar, void, bead, and cell regions (n = 18, 11, 28, and 84, respectively). RMS_H_, root-mean-square height fluctuation per object; ⟨RMS_H_⟩, mean RMS_H_ across objects. <C.V>, average coefficient of variation. Each data point represents an individual object.

We then validated FLEXOM height measurements using live L1210 cells stably expressing LifeAct–RFP, a genetically encoded F-actin marker that delineates the cell boundary [36]. Dual-channel imaging enabled co-registration of FLEXOM negative-contrast images with LifeAct–RFP-defined cell geometry from the same field of view (Figure 2d). FLEXOM-measured heights closely matched the confocal z-stack-derived reference profiles scaled to each cell dimension (Figure 2e; see Methods), with a linear correlation whose slope was statistically indistinguishable from unity (y = 1.05x − 0.77, R^2^ = 0.857; Figure 2f). This agreement validates the FLEXOM calibration scheme and its ability to measure absolute local cell height across the accessible height range.

We next quantified measurement precision using the per-object height fluctuation (See Methods). Silica bead height traces remained nearly flat over time (Figure 2g). Extending this analysis across platform references, beads, and live HUVEC cells showed that PDMS pillars and void regions defined the instrumental noise floor, whereas beads and cells exhibited sub-micrometer height fluctuations (Figure 2h; ⟨RMS_H_⟩ = 0.041 µm for pillars, 0.176 µm for voids, 0.366 µm for beads, and 0.325 µm for cells). Although live cells exhibited ⟨RMS_H_⟩ values comparable to those of beads, their larger inter-object dispersion indicated biological variability beyond measurement noise (Figure 2h; ⟨C.V.⟩ = 0.204 for cells vs. 0.050 for beads).

Finally, we asked whether the measurements remained stable during multi-day tissue monolayer development. Continuous imaging of A549 cells from sparse to fully confluent states over 65 hours yielded well-calibrated height maps throughout (Figure 3a and Video S1), with no detectable drift in the frame-by-frame intensity references. Together, these results validate FLEXOM as an accurate, sub-micrometer-precise, and stable platform for multi-day 3D topographic measurement of developing tissue monolayers.

**FIGURE 3.**
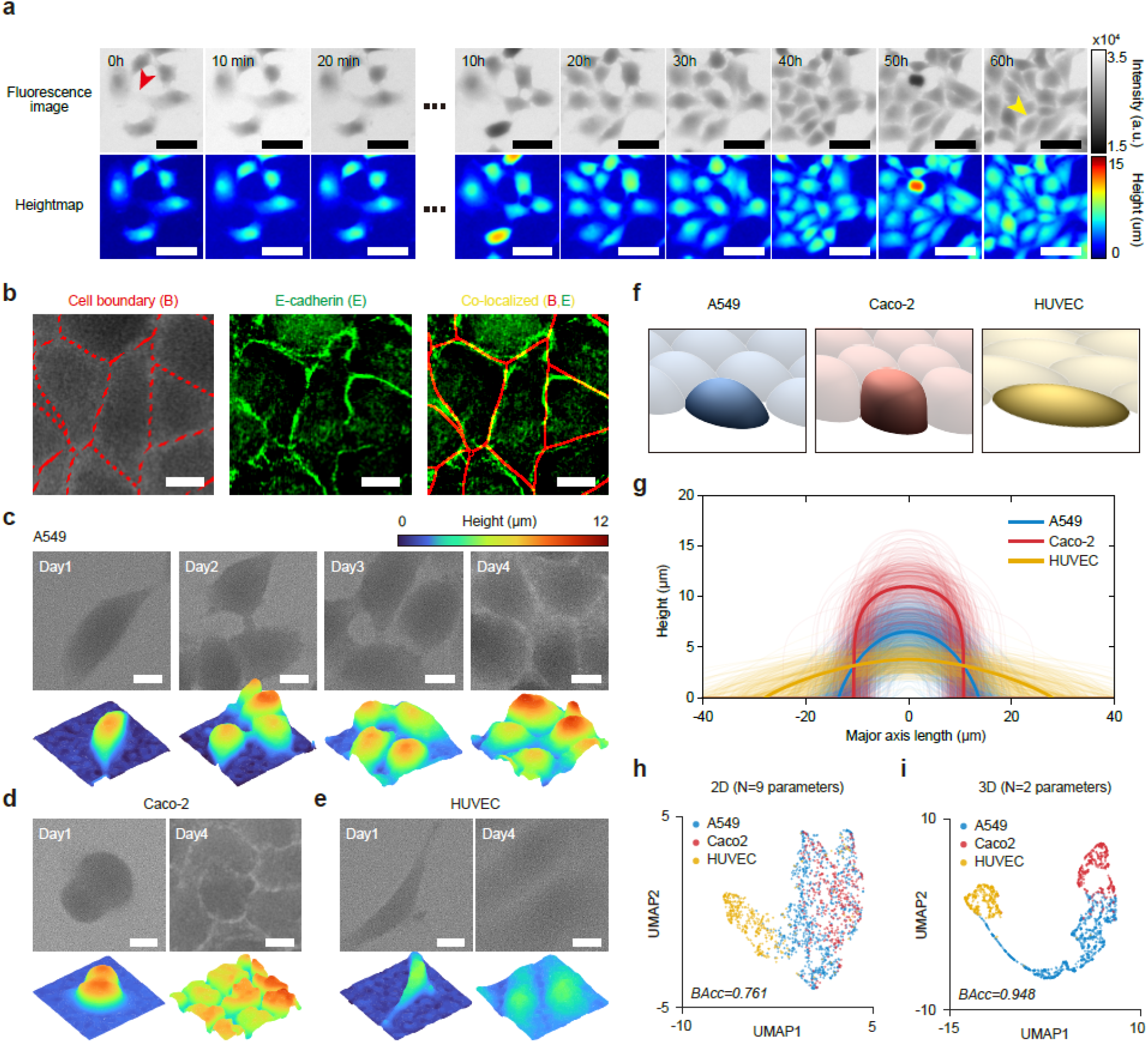
Vertical architecture as a structural signature of confluent tissue monolayers (a) Time-lapse fluorescence images (top) and corresponding 3D height maps (bottom) of A549 cells captured from sparse (0 h) to tissue monolayer formation (60 h) at 10 min intervals. Diffuse cell boundaries in the sparse regime (red arrowhead) and bright junctional lines upon confluence (yellow arrowhead) are indicated. Scale bars, 50 μm. (b) FLEXOM-derived cell boundaries (red) and E-cadherin-labeled cell–cell junctions (green); merged image shows co-localization (yellow). Scale bars, 20 µm. (c-e) Time-lapse 3D height maps of tissue monolayer formation from isolated cells (Day 1 through Day 4) for A549 (c), Caco-2 (d), and HUVEC (e) cells. Scale bars, 20 µm. (f) Representative 3D morphological renderings of individual cells reconstructed from group-averaged parameters (A549, n = 885 cells; Caco-2, n = 406 cells; HUVEC, n = 375 cells). (g) Cross-sectional height profiles of individual cells (thin lines) and per-cell-type mean profiles (thick lines) for the three cell types at confluence, corresponding to the representative in (f). (h,i) UMAP projections of single-cell morphologies based on nine 2D shape descriptors (area, aspect ratio, circularity, solidity, eccentricity, perimeter, extent, major axis, and minor axis; h) and two height-derived descriptors (mean cell height and height standard deviation; i). Balanced accuracy (BAcc) of cell-type classification using each parameter set is shown in the corresponding panel.

### 2.3 Vertical structure as an architectural descriptor of tissue monolayers

A tissue monolayer is assembled from individual cells that act as discrete structural units. Its overall topographic architecture is determined by two complementary features: the three-dimensional geometry of each cellular unit and the in-plane organization of these units, including their packing, spacing, and orientation. The in-plane component can be readily characterized using conventional 2D morphometric descriptors, including projected area and aspect ratio, as well as computational packing or tiling schemes such as Voronoi tessellation. By contrast, the vertical architecture of each cellular unit remains largely inaccessible to planar morphometry. We therefore used FLEXOM to resolve tissue monolayer topography at the single-cell level and extract cell-resolved 3D architectural signatures (Figure 3).

Single-cell topographic analysis requires delineation of individual cell boundaries within the confluent monolayer. Time-lapse imaging of A549 cells transitioning from sparse to confluent states revealed a pronounced evolution of the negative-contrast signal at cell boundaries (Figure 3a, Video S1). Isolated cells exhibited diffuse boundaries, whereas cells in tight cell–cell contact displayed sharp bright lines along their junctional interfaces, reflecting dextran accumulation in the narrow intercellular cleft. This emergent junctional contrast, a direct optical consequence of junctional network maturation, enabled label-free single-cell segmentation with the Cellpose-SAM model [37–39]. The resulting FLEXOM-derived boundaries co-localized closely with E-cadherin, a canonical marker of epithelial cell–cell junctions, in fixed confluent monolayers, confirming that the segmentation tracks the underlying junctional network and yields reliable single-cell masks for downstream topographic analysis (Figure 3b, Figure S5).

Applying this segmentation strategy, we quantified the evolution of 3D cellular topography in HUVEC, A549, and Caco-2 monolayers as they transitioned from isolated cells to compact monolayers with mature junctions (Figure 3c–e). Cell-type-specific differences in vertical morphology became markedly amplified across all pairwise comparisons upon monolayer formation (Figure 3c–f, Note S1, and Figure S6). At confluence, we observed nearly fivefold differences in average cell height across the cell types (Figure 3g; A549: 3.79 ± 0.06; Caco-2: 8.91 ± 0.11; HUVEC: 2.06 ± 0.04 μm).

We next tested whether vertical architectural signatures could distinguish individual cellular units from different monolayers more effectively than conventional planar descriptors. We projected single-cell morphological features into low-dimensional UMAP (Uniform Manifold Approximation and Projection) spaces and evaluated cell-type classification performance at the single-cell level. Notably, the vertical architecture-based representation outperformed the lateral-only representation by a large margin, despite using roughly fivefold fewer features; just two height-derived parameters—mean height and height standard deviation—achieved a classification accuracy of ~94.8%, outperforming models based on nine 2D parameters (e.g., area, circularity, solidity, and perimeter), which achieved an accuracy of ~76.1% (Figure 3h,i, Figure S7, and Note S1). These results show that the vertical geometry of individual cellular units is a primary structural signature across tissue monolayer types, revealing single-cell architectural differences that are largely missed by conventional planar descriptors.

### 2.4 Vertically anisotropic remodeling of tissue monolayers upon osmotic shock

Unlike passive materials, tissue monolayers actively adjust their 3D architecture in response to external perturbations. Osmotic response is a prime example of adaptive volume regulation [40–43]. Cells undergo rapid swelling or shrinking immediately after changes in extracellular osmolarity, followed by partial volume recovery through regulatory volume decrease or increase (RVD or RVI) [40,41,45]. Although osmotic volume regulation has been extensively studied in isolated single cells [42–44], it remains unclear how the same process occurs in densely packed monolayers, where neighboring cells constrain lateral deformation. Using FLEXOM, we tracked how osmotic shock remodels monolayer architecture along the vertical and lateral axes over time (Figure 4).

**FIGURE 4.**
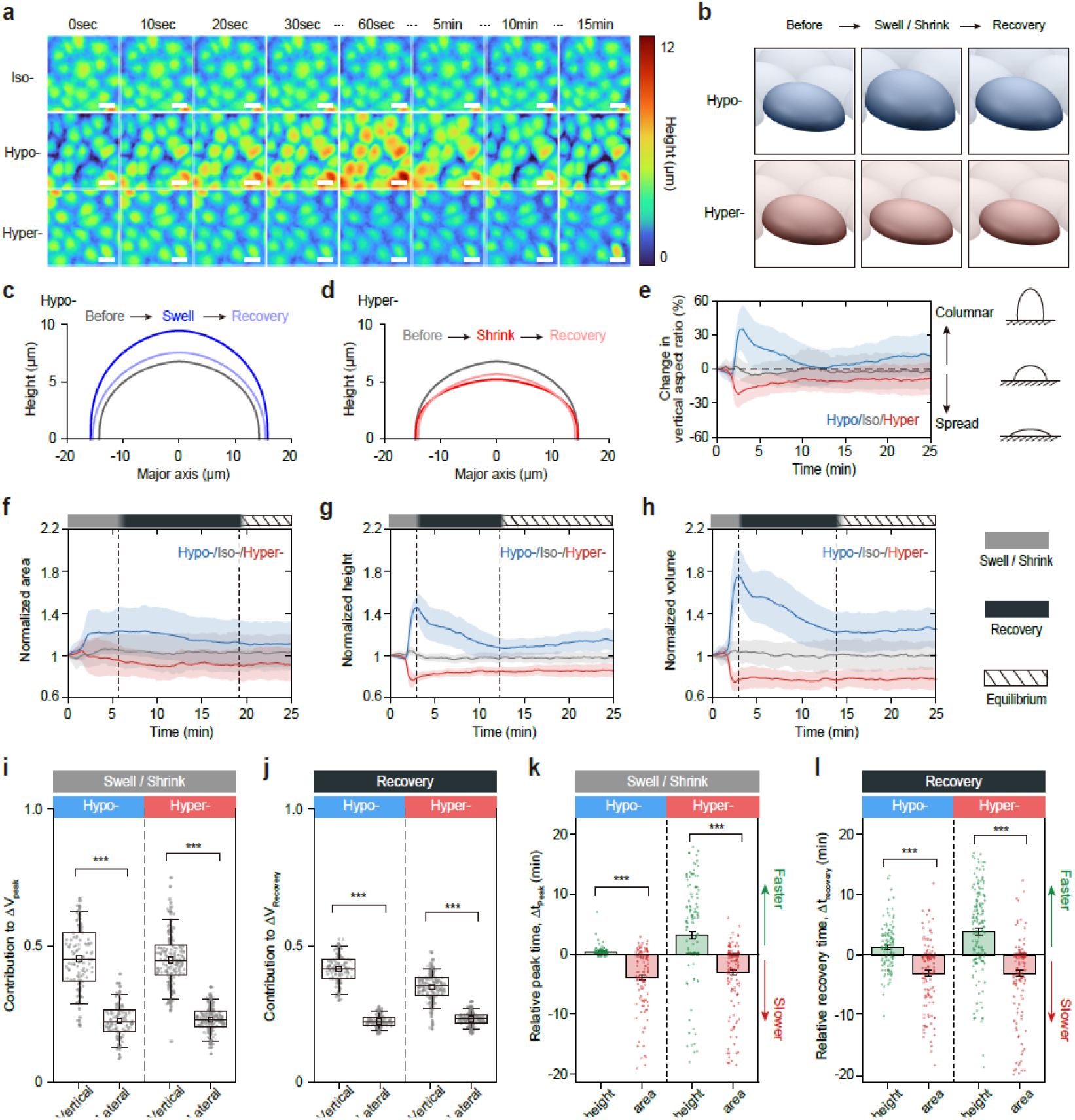
Anisotropic, height-driven osmotic volume regulation in tissue monolayers. (a) Time-lapse 3D height maps of A549 tissue monolayers under hypotonic (Hypo–; 150 mOsm L^−1^), isotonic (Iso–; 330 mOsm L^−1^), and hypertonic (Hyper–; 450 mOsm L^−1^) conditions. Scale bars, 20 µm. (b) Representative 3D renderings of individual cells before, during, and after osmotic perturbation, illustrating the characteristic swell–recovery and shrink–recovery responses. (c,d) Cross-sectional height profiles under hypotonic (c) and hypertonic (d) shock. (e) Temporal evolution of the normalized vertical aspect ratio (VAR ≡ mean height × area^−1/2^) under Hypo-(blue), Iso-(gray), and Hyper-(red) conditions; mean ± s.d. Inset, schematic cross-sections of columnar and spread tissue architectures. (f–h) Normalized projected area (f), mean cell height (g), and cell volume (h) over time under Hypo-(blue), Iso-(gray), and Hyper-(red) conditions; mean ± s.d. Color bars denote swell/shrink (gray), recovery (black), and equilibrium (striped) phases. Dashed lines mark osmotic phase transitions. (i,j) Vertical and lateral contributions to volume change during swell/shrink (i) and recovery (j) phases (n = 120 and 210 cells for Hypo- and Hyper-conditions, respectively). Box plots denote the interquartile range (box hinges), means (squares), and 5th/95th percentiles (whiskers). (k,l) Relative peak (Δt_peak_; k) and recovery (Δt_recovery_; l) times of mean height and projected area with respect to volume (n = 116 and 152 cells for Hypo- and Hyper-conditions, respectively). Δt = t_volume_ − t_feature_, with positive values denoting earlier response than volume and negative values denoting later response. Data depict mean (colored bars) ± s.d. (error bars). Data in (b–h), N = 3 (Hypo- and Hyper-) or 2 (Iso-) independent experiments, with n =120, 210, and 108 cells, respectively. Data in (i–l), statistical test: two-sided paired t-test. ***P < 0.001.

Confluent A549 monolayers (cultured for >4 days; Methods) were subjected to persistent transitions from isotonic (Iso–; 330 mOsm L^−1^) to either hypotonic (Hypo–; 150 mOsm L^−1^) or hypertonic (Hyper–; 450 mOsm L^−1^) conditions, and their 3D architecture was tracked at 10– second intervals (Figure 4a and Video S2). Under sustained osmotic perturbations (> 25 min), the 3D architecture of the tissue monolayer exhibited a characteristic two-step response comprising an initial deformation followed by partial recovery (Figure 4b), consistent with previous reports on RVI and RVD (Note S2). Interestingly, osmolarity-driven deformation of the tissue monolayer was not isotropic but was significantly anisotropic, biased toward the vertical axis (Figure 4c,d). Cross-sectional height profiles revealed that cells swelled and shrank primarily along the vertical axis, with comparatively small changes in lateral dimension under hypotonic and hypertonic shock, respectively. Consistently, the vertical-to-lateral aspect ratio increased by 35.6 ± 1.9% under hypotonic shock and decreased by 22.0 ± 0.9% under hypertonic shock at peak, followed by partial recovery to +11.3 ± 1.6% and −7.9 ± 1.0% relative to baseline (Figure 4e).

We next asked whether osmotic responses in tissue monolayers are driven primarily by vertical topographic modulation rather than lateral changes. In FLEXOM, cell volume is directly computed as projected area multiplied by mean height (Note S1), allowing volume changes to be partitioned into height-driven and area-driven contributions. To quantify this partitioning directly, we decomposed the instantaneous volume change along the vertical and lateral dimensions at each time step across the osmotic response. Strikingly, cell height contributed nearly twofold more than the lateral equivalent length to volume change, both at the peak response and throughout the recovery phase (Hypo–: C_vertical_ = 0.459 ± 0.010 and 0.419 ± 0.005 vs. C_lateral_ = 0.220 ± 0.005 and 0.219 ± 0.002, respectively; Hyper–: C_vertical_ = 0.450 ± 0.006 and 0.349 ± 0.003 vs. C_lateral_ = 0.229 ± 0.003 and 0.230 ± 0.002, respectively; Figure 4i,j and Note S2). Vertical remodeling therefore outweighs lateral-area adjustments as the dominant geometric contributor to osmotic volume change in confluent tissue monolayers.

Finally, we investigated the temporal decoupling between the vertical and lateral architectural dynamics. Under both hypotonic and hypertonic perturbations, vertical height and volume shifted abruptly following the shock, whereas the projected area displayed a gradual drift across a broader temporal window (Figure 4f-h). Height reached its peak earlier than volume, while the projected area reached its extremum well after volume under both osmotic conditions (Hypo–: Δt_peak, height_ = 0.4 ± 0.1 min, Δt_peak, area_ = −3.9 ± 0.5 min; Hyper–: Δt_peak, height_ = 3.1 ± 0.6 min, Δt_peak, area_ = −3.1 ± 0.4 min; Figure 4k and Note S2). A similar temporal decoupling was observed during recovery: height relaxation preceded volume stabilization, whereas the projected area exhibited pronounced delays relative to volume under both perturbations (Hypo–: Δt_recovery, height_ = 1.3 ± 0.4 min, Δt_recovery, area_ = −3.0 ± 0.5 min; Hyper–: Δt_recovery, height_ = 3.9 ± 0.6 min, Δt_recovery, area_ = −3.0 ± 0.5 min; Figure 4l and Note S2). Consistently, time-series correlation analysis showed that height dynamics track volume dynamics more closely than the projected area (Figure S8), indicating that height is a better proxy of volume homeostasis during tissue monolayer’s osmoregulation.

### 2.5 Axially dominated response under cyclic osmotic driving

Because lateral remodeling was several minutes slower than vertical remodeling under sustained osmotic shock, we hypothesized that rapid cyclic stimulation could temporally separate the vertical and lateral components of the response. Specifically, cycling the osmotic input faster than this temporal delay would suppress lateral remodeling while preserving rapid vertical deformation, therefore accommodating volume oscillation entirely along the vertical axis.

To test this hypothesis, we subjected confluent A549 monolayers to periodic switching between isotonic (330 mOsm L^−1^) and hypotonic (150 mOsm L^−1^) conditions at a 3 min alternation period, tracking the 3D architecture at 250 ms per frame over 30 min (Figure 5a). Upon each hypotonic pulse, the monolayer swelled rapidly within seconds. Before the delayed, active osmotic shock response (i.e. RVD) could be initiated, we quickly induced deswelling by switching the monolayers back to isotonic solution. As a result, cell volume within the monolayer rose and fell in close synchrony with the osmotic waveform, and cell height closely followed this trajectory, whereas projected area remained nearly unchanged across cycles (Figure 5b–d). Quantitative comparison of isotonic and hypotonic phases confirmed this pattern statistically: volume and height increased during hypotonic phases, whereas projected area did not change significantly (Figure 5e–g, volume, 1529 ± 2 vs. 2085 ± 3 μm^3^, P < 10^−4^; height, 2.793 ± 0.002 vs. 3.814 ± 0.005 μm, P < 10^−4^; area, 558.4 ± 0.6 vs. 556.0 ± 0.6 μm^2^, P = 0.669; isotonic vs. hypotonic). Thus, during cyclic osmotic stimulation, the monolayer remodeled mainly along the vertical axis, while its lateral architecture remained nearly unchanged throughout (Figure 5h). Together, these results establish stimulus timing as an independent control parameter for directional tissue remodeling, separating fast vertical deformation from slower lateral remodeling without changing stimulus magnitude.

**FIGURE 5.**
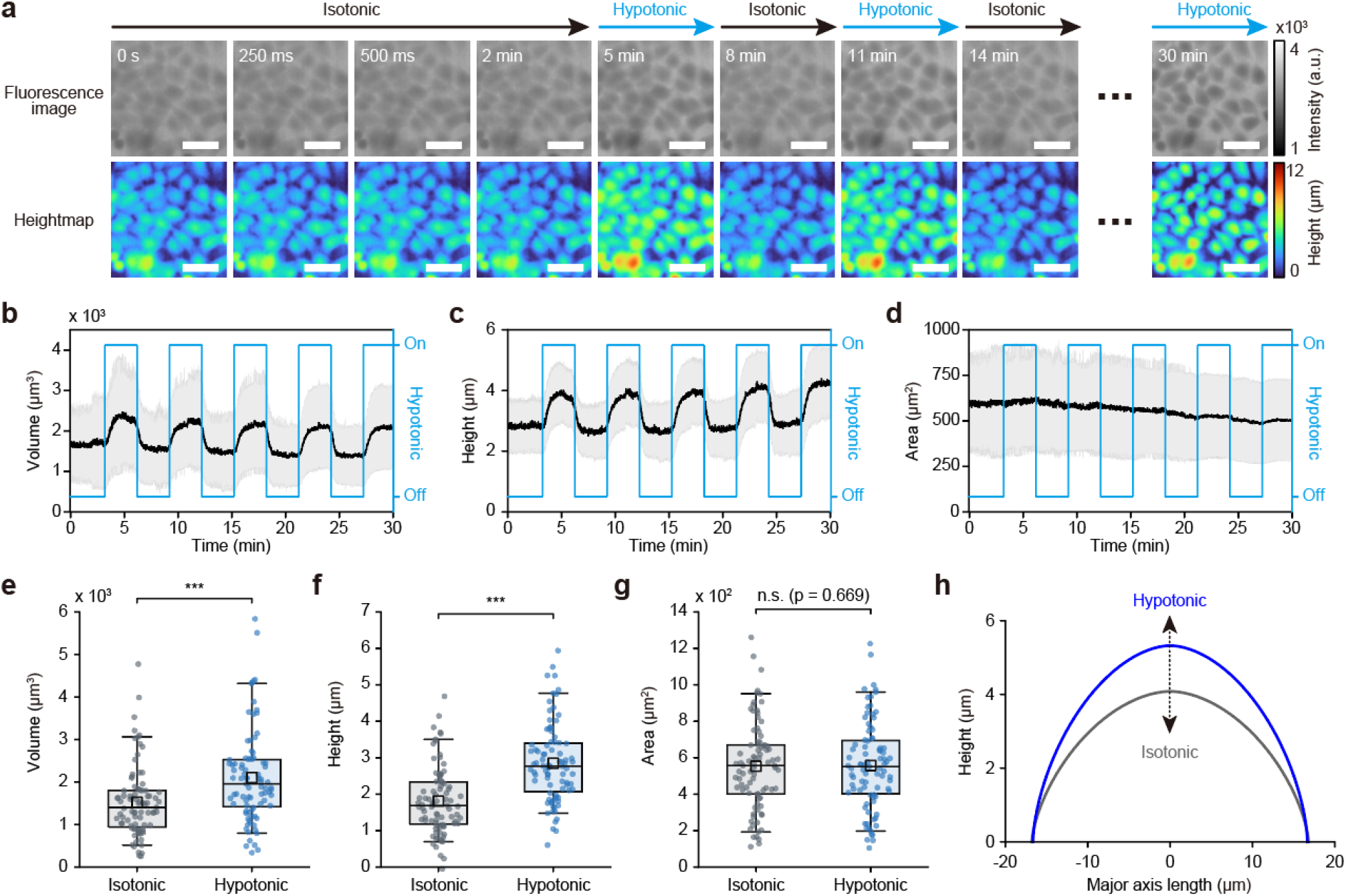
Axially dominated response of tissue monolayers under rapid and cyclic osmotic switching. (a) Time-lapse fluorescence images (top) and corresponding 3D height maps (bottom) of an A549 tissue monolayer subjected to cyclic osmotic perturbations over 30 min at 250 ms per frame. After an initial 2 min isotonic baseline (black arrow; 330 mOsm L^−1^), hypotonic (blue arrow; 150 mOsm L^−1^) and isotonic solutions were alternated every 3 min. Scale bars, 50 μm. (b–d) Cell volume (b), projected area (c), and mean height (d) of individual A549 cells within the same tissue monolayer (n = 87 cells from N = 3 regions of interest). Right axes (blue) indicate the osmotic stimulation state (On, hypotonic; Off, isotonic). Data are presented as mean (black line) ± s.d. (gray shading). (e–g) Phase-wise comparisons of cell volume (e), mean height (f), and projected area (g) between isotonic (gray) and hypotonic (blue) phases during cyclic osmotic switching. Each point represents the mean value at a single time point from the corresponding traces in (b–d). Box plots denote the interquartile range (box hinges), means (squares), and 5th/95th percentiles (whiskers). n.s., not significant; ***, P < 10^−4^. (h) Phase-averaged cross-sectional height profiles during rapid cyclic osmotic stimulation (isotonic, gray; hypotonic, blue).

## 3. Discussion

Here, we introduce FLEXOM, which provides frame-by-frame 3D topographic profiling of confluent tissue monolayers with self-calibrated, label-free operation on a standard wide-field microscope (Figure 1). Utilizing the rapid profiling rate (~250 ms per frame), sub-micrometer axial precision, and multi-day operational stability (>96 h; Figure 2) of FLEXOM, we captured a wide variety of 3D architectural and dynamical features across cell types and perturbation regimes that were previously unknown.

One notable finding is that height-derived metrics outperformed a larger set of 2D shape metrics in single-cell classification (Figure 3), indicating that vertical architecture captures structural variation missed by conventional lateral morphometry. One interpretation is that cell height reflects mechanical variables that are poorly represented in planar shape metrics, including cortical tension, pressure, adhesion, and junctional mechanics. This view is consistent with prior studies [47,48] showing that epithelial height and 3D shape are controlled by adhesion and actomyosin contractility, and that epithelial sheets can exhibit material-like tension–strain behavior in three dimensions. These findings also carry significant implications for tissue modeling. Widely used vertex, Voronoi, and jamming models describe confluent monolayers as planar tilings where mechanics and dynamics are encoded via 2D quantities (e.g., cell area, edge tension, T1 transition frequency) [18,19,49–51]. Because cell height varies strongly across cell types, 2D model fits may misattribute height-dependent variation to effective planar parameters, potentially masking true mechanical differences between cell types or tissue states. Could vertical topography make mechanical inference in tissue models more reliable? Integrating FLEXOM-measured height fields into 3D vertex or active-surface models [48,51,52] could improve mechanical inference by helping distinguish true mechanical variation from parameter compensation caused by missing 3D structure.

Next, we observed that osmotic shock-induced anisotropic responses occur in both sparse cells (Figure S9) and confluent monolayers (Figure 4). In fully packed monolayers, mature cell-cell junctions may provide lateral constraints that direct osmotic deformation preferentially along the vertical axis. However, the persistence of height-dominated deformation even in sparse cells indicates that junctional confinement alone cannot fully account for this anisotropy. Another potential contributor is cell–substrate adhesion, which transmits actomyosin-generated traction forces to the substrate via the basal interface [6,53]. Since our projected area measurements reflect the lateral cellular footprint, basal adhesion is likely a major determinant of lateral deformation at the single-cell level. Specifically, A549 cells have been reported to transmit basal traction forces of 55.0 ± 12.0 nN [54], comparable to the osmotic deformation force on the order of 100 nN, estimated from reported rounding pressure of ~140 Pa [55], scaled by our measured cell surface area. These suggest that basal traction provides resistance to lateral osmotic deformation, thereby providing a simple mechanical explanation for the approximately twofold bias toward vertical remodeling.

By contrast, the height–area temporal decoupling was substantially weaker in sparse cells (Figure S9). While osmotic remodeling primarily involves intracellular cytoskeletal force balance in isolated cells, the shared cell–cell junctions in compact monolayers may act as additional mechanical damping elements [56–58]. Deformation of these junctional boundaries perturbs junctional tension and transmits force changes to neighboring cells [57,59]; thus, lateral remodeling requires global force rebalancing across the monolayer that extends beyond the individual intracellular cytoskeletons. Notably, the minute time scale height-area temporal delay we observed is consistent with reported mechanical stress-relaxation times in epithelial monolayers [60]. A second source of height–area temporal decoupling may be the active turnover of junctional components. E-cadherin, a core component of adherens junctions, is continuously recycled and reassembled at mature cell–cell contacts on a minute timescale [61]. Collectively, these intercellular rebalancing steps and biochemical turnover may explain why lateral remodeling is delayed in compact monolayers relative to sparse cells.

Our work also suggests that temporal pattern of the stimulus, independent of its magnitude, can tune tissue architectural responses along specific geometric axes. In particular, minute-period cyclic osmotic perturbation confined tissue deformation exclusively to the vertical axis (Figure 5). This suppression of lateral remodeling may arise from two distinct mechanisms: (i) lateral remodeling is an active response downstream of canonical osmotic response cascades such as RVD or RVI; or (ii) lateral deformation is a passive response that requires slow mechanical force relaxation before becoming evident. In the first scenario, lateral remodeling may depend on active, time-intensive cellular processes such as osmolyte transport, RVD/RVI-associated signaling, actomyosin remodeling, and adhesion or junctional reorganization [40,41,45,46]. Rapid switching to isotonic condition could therefore ‘reset’ the osmotic state before this active lateral remodeling is initiated. In the second scenario, passive osmotic water flux may immediately initiate lateral deformation [40-42], but mechanical dampers including basal adhesion and junctional confinement simply delay lateral expression beyond the perturbation period. These hypotheses could be tested by perturbing volume-regulatory pathways such as VRAC/LRRC8 [45]. More broadly, FLEXOM may complement and initiate studies that quantify the temporal dynamics of intracellular signaling cascades involved in RVD or RVI, and determine whether these pathways merely correlate with, or actively drive 3D mechanical anisotropy in living matter.

FLEXOM nonetheless has several technical limitations (Note S3). First, FLEXOM reports the total thickness of the dye-excluding region. For adherent cells, this represents the height profile; however, for suspended cells, a spherical cell may appear as a hemisphere resting on the floor. Second, FLEXOM may misattribute thickness when cells overlap along the vertical axis, for instance during apical extrusion, when one cell rides over another, and ceiling attachment. In such cases, FLEXOM reports the combined thickness, which can falsely suggest that the underlying cell is taller. Third, there is a trade-off between dynamic range and axial resolution set by the microfluidic channel height. Finally, FLEXOM may be particularly sensitive to refractive-index mismatch between the medium and the sample as imaging specimens become thicker. We anticipate that combining FLEXOM with high–numerical aperture (NA) imaging will potentially resolve these limitations. At higher NA, fluorescence collection becomes more axially weighted, with signal preferentially arising from planes near the focal region. This could help distinguish thickness contributions from different axial layers, reducing z-position ambiguity for suspended geometries and improving interpretation in overlapping-cell configurations. Future work combining low- and high-NA imaging, together with a computational framework to infer the axial origin of fluorescence, may expand FLEXOM application to thicker samples, such as multilayers and organoids.

## 4. Experimental Section

### 4.1 Micropatterned master fabrication

SiO2 wafers were cleaned by SPM (H2SO4:H2O2=3:1), rinsed with DI water, and dehydrated. SU-8 2025 negative photoresist (KAYAKU) was spin-coated at 500 rpm for 10 s then 3,000 rpm for 30 s to cover ~60% of the wafer surface, and left at room temperature for ~24 h to allow planarization. Soft bake was performed on a leveled hotplate at 65 °C for 3 min then 95 °C for 5 min. Samples were exposed on a MA6 i-line aligner (365 nm) at 25 mW cm^−2^ for 6 s (total dose: 150 mJ cm^−2^), followed by post-exposure bake at 65 °C for 1.5 min then 95 °C for 5 min. Wafers were developed in SU-8 developer (KAYAKU) for 5 min with gentle agitation, rinsed sequentially in IPA (1 min) and DI water, dried with N2 gun, and hard-baked at 200 °C for 10 min.

### 4.2 Microfluidic chip fabrication

Polydimethylsiloxane (PDMS; Sylgard 184, Dow Corning) was mixed with a curing agent at a 10:1 (base:agent) ratio and degassed under vacuum for 30–60 min. The mixture was poured over the SU-8 patterned master and vacuum-treated for 2 h to ensure complete pattern filling. PDMS slabs (~4 mm thick) were cured at 80 °C for 2 h, peeled from the master, and inlet/outlet ports were punched using a 0.5 mm diameter biopsy punch. The PDMS surface was cleaned with adhesive tape and bonded to a glass coverslip by oxygen plasma treatment (50 W, 30 s) to form a closed microfluidic chamber. After plasma bonding, the assembled device was placed in an 80 °C oven for 15 min to reinforce bonding, then stored overnight at room temperature to allow surface hydrophobic recovery. The following day, 70% ethanol was applied to both inlet and outlet ports, and PEEK tubing (1/16″OD × 0.007″ID; IDEX, 1536L) was gently inserted into the 0.5 mm punched holes for fluidic connection. For long-term culture and imaging experiments, the chip was interfaced with a custom pneumatic control system. Fluid reservoirs (glass vials; Wheaton) pressurized with 5% CO2-regulated gas were connected to the chip via the PEEK tubing. Flow was driven by electronic pressure regulators (ITV1100, SMC) and miniature solenoid valves (S070, SMC), controlled via National Instruments I/O hardware and a custom LabVIEW 2019 routine synchronized to image acquisition.

### 4.3 FLEXOM imaging systems

All imaging was performed on an inverted fluorescence microscope (Ti2-E, Nikon Instruments) equipped with a motorized XYZ stage and a scientific sCMOS camera (Zyla 5.5, Andor). A Plan Apo λ 20×/0.75 NA objective (Nikon) was used for all experiments unless otherwise noted. The microfluidic chip was mounted on a temperature-controlled stage (LCI) maintained at 37 °C. Time-lapse FLEXOM imaging was conducted using a GFP filter set (excitation 475 nm, emission 525 nm) with 2,000 kDa FITC–dextran (1 mg ml^−1^; FD2000S, Sigma-Aldrich) dissolved in the culture medium as the exclusion dye. Unless otherwise stated, Exposure time (25 ms) and illumination intensity (10 %) were kept constant across all experiments to ensure quantitative consistency. All analyses were performed on single-view images without stitching.

### 4.4 Cell culture

A549 and Caco-2 cells were kindly provided by Dr. Junsang Doh (Seoul National University) and maintained in Dulbecco’s Modified Eagle Medium (DMEM; Gibco) supplemented with 10% fetal bovine serum (FBS; Gibco) and 1% penicillin–streptomycin at 37 °C in a humidified 5% CO2 incubator. Human umbilical vein endothelial cells (HUVEC; ScienCell, Catalog #8000) were cultured in Endothelial Cell Medium (ECM; ScienCell, Catalog #1001) supplemented with the complete kit provided by the manufacturer. All media were replaced every 2–3 days, and cells were passaged using 0.25% trypsin–EDTA before reaching 90% confluency. A549, Caco-2, and HUVEC cells were tested for mycoplasma contamination, and no contamination was detected.

### 4.5 In-chip cell seeding and culture

Microfluidic chips were sterilized sequentially by infusing 70% ethanol, DI water, phosphate-buffered saline (PBS), and complete culture medium for at least 5 min each. From the washing step onward, chips were maintained in a 37 °C, 5% CO2 incubator to equilibrate temperature and gas composition, thereby maintaining medium pH, given the gas permeability of PDMS. Cell suspensions were loaded into the microfluidic chamber from pressurized glass vials at ΔP ≈ 100 mbar for 1 min, then allowed to attach under static conditions for 4 h. During subsequent culture, the device was kept static between acquisitions and briefly perfused (ΔP ≈ 50 mbar, 1 min every 10 min) to replenish fresh medium while minimizing shear stress. For confluent monolayer experiments, cells were seeded at ~8 × 10^6^ cells ml^−1^ and cultured for >4 days to ensure formation of compact monolayers with mature cell–cell junctions prior to measurements. For sparse cell experiments, cells were seeded at ~1 × 10^6^ cells ml^−1^ and imaged within 24 h of seeding to ensure non-confluent conditions.

### 4.6 Evaluation of FITC–dextran uptake and biocompatibility

To assess size-dependent cellular uptake of dextran, cells were seeded into 35 mm glass-bottom confocal dishes and grown to 90–100% confluency (~3 days post-seeding). Culture medium was replaced with fresh medium containing 1 mg ml^−1^ of either 2,000 kDa FITC–dextran (FD2000S, Sigma-Aldrich) or 70 kDa TRITC–dextran (D1819, Invitrogen) and incubated for 24 h. Nuclei were counterstained with Hoechst 33342 (10 μg ml^−1^) for 15 min, followed by two washes with fresh medium to remove extracellular dye. FITC–dextran and TRITC–dextran were imaged using GFP (Ex/Em: 475/525 nm) and Cy3 (Ex/Em: 550/605 nm) filter sets, respectively.

To assess biocompatibility, cells were seeded at 1 × 10^5^ cells ml^−1^ in 24-well plates and cultured in a medium supplemented with 2,000 kDa FITC–dextran (1 mg ml^−1^) or regular medium as a control. Cell proliferation was monitored by time-lapse phase-contrast imaging at 1 h intervals for 3 days using Incucyte SX5 (NFEC-2025-08-307698, Sartorius) with a 10× objective.

### 4.7 Image processing and height calibration

Raw fluorescence images were preprocessed in NIS-Elements (Nikon) as follows. First, shading correction was applied to compensate for non-uniform illumination (vignetting) using a reference image acquired from a pillar-free region within the microfluidic chamber representing uniform fluorescence at maximum chamber height (*H*_*max*_). Second, a 2D Fast Deconvolution algorithm was applied to enhance cell boundary contrast and suppress out-of-focus haze. Third, a 5 × 5 spatial median filter (medfilt2) was applied to remove single-pixel outliers such as hot pixels. The microchamber height *H*_*max*_ was experimentally determined to be 27 μm by profilometry of the SU-8 master mold (Alpha-Step D-500, KLA-Tencor).

Pixel fluorescence intensity I(*x,y,t*) was converted to quantitative cell height h(*x,y,t*) based on the fluorescence-exclusion principle [32], using frame-specific calibration references extracted from the multilayered micropillar array:

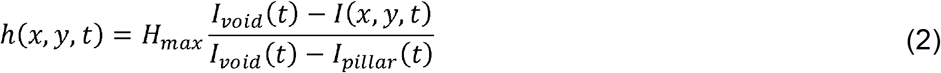

where *I*_*pillar*_(*t*) and *I*_*void*_(*t*) were extracted from manually selected rectangular reference ROIs placed over full-height PDMS pillar regions and enclosed cell-free void regions, respectively. *I*_*pillar*_(*t*) was defined as the minimum intensity within the pillar ROI, corresponding to *H*_*max*_, whereas *I*_*void*_(*t*) was defined as the maximum intensity within the void ROI, corresponding to h = 0. Both calibration references were re-evaluated for every frame to compensate for excitation drift and photobleaching. To account for spatial intensity variations between the interior of the multi-micropillar array and the surrounding open region, a position-specific correction factor was derived from the initial frame and applied to *I*_*void*_(*t*) throughout the time-lapse sequence. Pixels with calculated heights outside the physical range (h < 0 or h > *H*_*max*_) were clamped to 0 and *H*_*max*_, respectively. The lateral pixel resolution was 0.325 μm pixel^−1^ and the axial dynamic range spanned 27 μm over a 16-bit intensity range. The resulting calibrated height maps were used for all subsequent single-cell morphometric and volumetric analyses.

### 4.8 Accuracy validation of height measurements

To validate height measurement accuracy, LifeAct-RFP L1210 cells were previously generated [36] by transducing L1210 cells with an F-actin-labeling lentiviral vector (rLVUbi-LifeAct-TagRFP, ibidi GmbH). These cells were loaded into the microfluidic chip and allowed to settle on the substrate. Reference 3D cell profiles were acquired by confocal laser scanning microscopy (LSM 700, Zeiss) using a C-Apochromat 40×/1.20 W Korr water-immersion objective. LifeAct–RFP signals were captured at 555 nm excitation with a PMT detector (585–800 nm emission window) and a pinhole of 0.97 Airy units. Reference axial profiles were generated by tracing cell boundaries from side-view (x–z) projections of confocal z-stacks.

To construct a normalized reference profile, binary masks from confocal x-z images were converted into one-dimensional thickness profiles by summing mask pixels along each lateral position (Figure S10). The lateral coordinate was centered at the cell centroid and normalized by the basal half-width r, and the thickness was normalized by the same length scale. The averaged dimensionless profile was then fitted to a symmetric dome model:

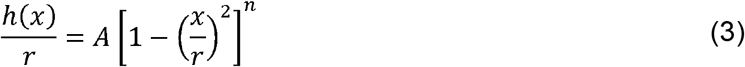

where *h(x)* is the local cell height, *x* is the lateral position from the centroid, *r* is the basal half-width, and *A* and *n* are fitting parameters defining the apex-to-base ratio and profile curvature, respectively.

In a separate FLEXOM measurement, LifeAct–RFP L1210 cells were imaged using GFP/Cy3 filter sets to simultaneously capture height maps and F-actin boundaries. For each FLEXOM-imaged cell, the fitted confocal-derived profile was scaled to the basal diameter measured from the LifeAct–RFP channel to generate a cell-specific modeled height profile. Cross-sectional FLEXOM-measured height profiles along the x- and y-axes through the cell centroid were then compared with the modeled profiles after pairing cells with closely matched basal diameters. Height measurement accuracy was assessed by linear regression between FLEXOM-measured and modeled height values, using only positions where the modeled height exceeded 0 μm

### 4.9 Precision quantification of height measurements

To quantify height measurement precision, silica beads (#904368; Sigma-Aldrich) and HUVEC cells were loaded into the microfluidic chip alongside pillar and void regions as structural controls. Mean height was calculated from a 5 × 5 pixel (0.325 μm pixel^−1^) ROI for each region type: pillar regions were sampled from areas not used for calibration; void regions were sampled from cell-free areas outside the multi-micropillar structure; and bead and cell regions were sampled from a 5 × 5 pixel window centered on individual beads or cells, respectively. The root-mean-square height fluctuation of each object or region was calculated as

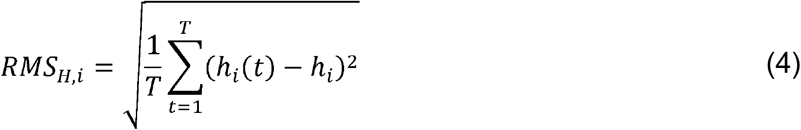

where *hi(t)* is the mean height of object or region *i* at time *t, hi* is its time-averaged height_*i* i_ is its time-averaged height, and *T* is the total number of frames. The ensemble-averaged precision was defined as

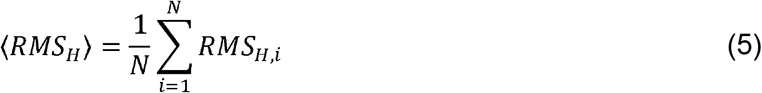

and inter-object variability was quantified by the coefficient of variation,

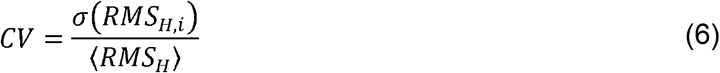

All measurements were performed at 1 min per frame over 25 min in a 27 μm-high chamber using a Plan Apo λ 20×/0.75 NA objective. The lateral pixel size was 0.325 μm pixel^−1^.

### 4.10 Junction staining and co-localization

To validate FLEXOM-derived cell boundary detection, cells cultured in the microfluidic chip for 4 days were first imaged by FLEXOM (GFP channel) to acquire fluorescence exclusion images of live monolayers. Cells were then fixed with 4% paraformaldehyde (PFA) in PBS for 10 min at room temperature, permeabilized with 0.2% Triton X-100 in PBS for 10 min, and blocked with 1% bovine serum albumin (BSA) in PBS for 1 h, with two PBS washes between each step. Cells were incubated overnight at 4 °C with a primary antibody against E-cadherin (1:200 dilution; #3195; Cell Signaling Technology), followed by three PBS washes and incubation with an Alexa Fluor-conjugated secondary antibody (1:500 dilution; #A11008, Invitrogen) for 2 h at room temperature protected from light. After three final PBS washes, E-cadherin immunofluorescence images were acquired using the same GFP channel. The micropillar array served as a fixed positional reference to align the pre- and post-fixation images in x-y. A cell boundary mask was generated from the live FLEXOM image using Cellpose, and boundary contours were extracted from the resulting label mask and overlaid onto the spatially aligned E-cadherin immunofluorescence image for qualitative comparison.

### 4.11 Cell segmentation and tracking

Cell boundaries were segmented from preprocessed FLEXOM fluorescence images using Cellpose (v4.0) [37] with the Cellpose-SAM pretrained model (CPSAM) [38], which incorporates a segment-anything model (SAM)-based architecture to improve boundary detection in densely packed monolayers [39]. Prior to segmentation, images were percentile-normalized by linear rescaling between the 1st and 99th intensity percentiles to standardize the input dynamic range across time. Segmentation was performed with the parameters flow_threshold = 0.4, cellprob_threshold = 0.0, and invert = True to account for the negative contrast inherent to fluorescence exclusion. The dynamics iteration parameter was set to niter = 10,000. Segmented masks were assigned unique per-cell labels and propagated across time frames using the TrackMate plugin in Fiji [62,63]. Segmented masks were processed using the Label Image Detector and linked via the Sparse LAP Tracker with the following parameters: maximum frame-to-frame linking distance of 30 pixels, maximum gap-closing distance of 50 pixels, and a maximum frame gap of 3 frames. Track splitting and merging were disabled. Final output images were generated by remapping pixel values of each cell mask to its corresponding unique Track ID to ensure consistent labeling across time.

### 4.12 Morphological feature extraction

Single-cell morphological features were quantified from reconstructed height maps and labeled segmentation masks using custom MATLAB (MathWorks) scripts. A 11–dimensional phenotypic feature vector was extracted for each cell, comprising nine 2D shape descriptors (projected area, perimeter, major and minor axis lengths, aspect ratio, circularity, solidity, eccentricity, and extent) and two height-derived topographic parameters (mean cell height and height standard deviation). For each segmented cell i occupying the projected domain Ωi, projected area and mean height were calculated as

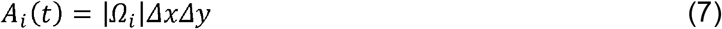

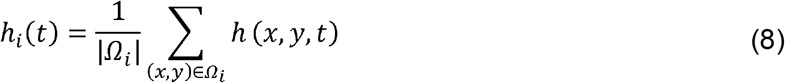

where Δ*x* = Δ*y* = 0.325 ◻ μm. Cell volume was then computed by integrating the calibrated height map over the cell footprint:

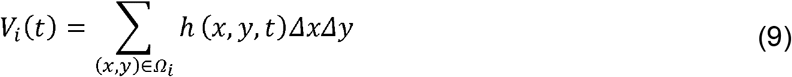

This is equivalent to V◻ (t) = A◻ (t) × *h*_*i*_ (*t*)when is defined as the footprint-averaged height, and was used for all osmotic-response analyses. See Figure S5 and Note S1 for additional detailswhen

### 4.13 Osmotic perturbation experiments

A549 monolayers were cultured in the chip for >4 days prior to imaging. A baseline medium (330 mOsm L^−1^) was prepared using DMEM supplemented with 10% FBS. Hypotonic (150 mOsm L^−1^) medium was prepared by diluting the baseline medium with DI water. Hypertonic (450 mOsm L^−1^) medium was prepared by supplementing the baseline medium with D-mannitol (Sigma-Aldrich). Isotonic control (330 mOsm L^−1^) medium was prepared by substituting DI water with PBS at an equivalent volume ratio, serving as a medium-exchange control. All media contained 1 mg ml^−1^ 2,000 kDa FITC–dextran. For sustained (unidirectional) perturbation experiments, monolayers were imaged during a 5 min baseline at 10 s per frame, followed by medium exchange to hypotonic, hypertonic, or isotonic perturbation conditions with continued imaging for 35 min. For cyclic (bidirectional) perturbation experiments, hypotonic and isotonic solutions were alternated every 3 min over 30 min following a 2 min baseline, with imaging at 250 ms per frame throughout. To account for experimental variability in tubing length, osmotic stimulation timing at the tissue was aligned to monolayer volume changes rather than to the programmed valve-switching time. The first abrupt change in monolayer volume was used to define the effective arrival of the stimulus at the tissue. In sustained perturbation experiments, the time point 2 min before this change was set as t = 0 min. In cyclic perturbation experiments, hypotonic and isotonic phases were assigned alternately at 3 min intervals from the effective stimulus arrival time.

### 4.14 Recursive ceiling-height calibration for cyclic osmotic perturbations

During cyclic bidirectional perfusion, changes in flow direction alter the local pressure distribution within the chamber, producing transient intensity drift in regions of interest (Figure S11). To correct this pressure-dependent calibration shift, recursive ceiling-height calibration was applied to the cyclic perturbation datasets (Figure S12). For each frame, the void and pillar reference intensities were first used to calculate a dimensionless height coefficient:

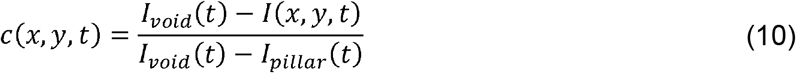

The local cell height was then reconstructed as

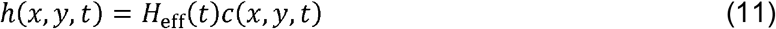

where *H*_*eff*_*(t)* is the effective chamber height used for height reconstruction at frame t. At each flow-direction switch, *H*_*eff*_*(t)* was recursively updated to preserve the pre-switch cell height after the pressure-induced intensity shift. The updated chamber height was estimated by least-squares fitting between the pre-switch cell height and the post-switch height coefficient:

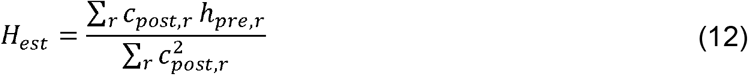

where *h*_*pre,r*_ is the pre-switch cell height and *c*_*post,r*_ is the post-switch height coefficient measured in region r. The estimated *H*_*est*_ was used as *H*_*est*_ for subsequent frames until the next flow-direction switch. Each update was computed from a 5-frame median around the switching point to suppress single-frame noise. Corrected height maps were used to extract single-cell morphology over time, and trajectories were smoothed with a centered 5-frame moving median before analysis.

### 4.15 Statistical analysis and software

Image preprocessing was performed in NIS-Elements (Nikon). Cell segmentation was performed using Cellpose (v4.0) with the Cellpose-SAM pretrained model (CPSAM). Cell tracking was performed using the TrackMate plugin in Fiji (ImageJ). All subsequent image processing and quantitative data analyses were performed using custom scripts written in MATLAB (MathWorks, R2023b) and Python (v3.10). Dimensionality reduction (UMAP) and cell-type classification (linear SVM with ECOC strategy) were implemented using the umap-learn Python package and scikit-learn library, respectively. Trajectory phase segmentation was implemented using the ruptures Python package (PELT algorithm). Statistical comparisons between two independent groups were performed using two-sided Welch’s t-test; paired comparisons within the same cells were performed using two-sided paired t-tests. Cell-type classification performance was evaluated by balanced accuracy (BAcc) using 5-fold stratified cross-validation (Note S1). Effect sizes were calculated using Cohen’s d. The null hypothesis was that mean values are equal across groups. All reported P values are two-sided. Data are presented as mean ± s.d. unless otherwise stated. Sample sizes (n, number of cells or objects; N, number of independent experiments or regions of interest) for each dataset are indicated in the figure legends.

### 4.16 Representative morphology reconstruction

See Note S1 for details.

### 4.17 Dimensionality reduction and classification accuracy

See Note S1 for details.

### 4.18 Analysis of vertical and lateral geometric contributions

See Note S2 for details.

### 4.19 Quantification of osmotic response dynamics

See Note S2 for details.

