## Supplementary Data for "Continuous, high-speed 3D topographic profiling of living tissue monolayers"

*

[**Note S2. Anisotropic and height-dominated volume regulation
during osmotic shock**	1](#_Toc228204412)7

[**Note S3. Measurement limitations and device constraints**](#_Toc228204413)	20

[**Supplementary References**	2](#_Toc228204409)2

**Supplementary Figures**

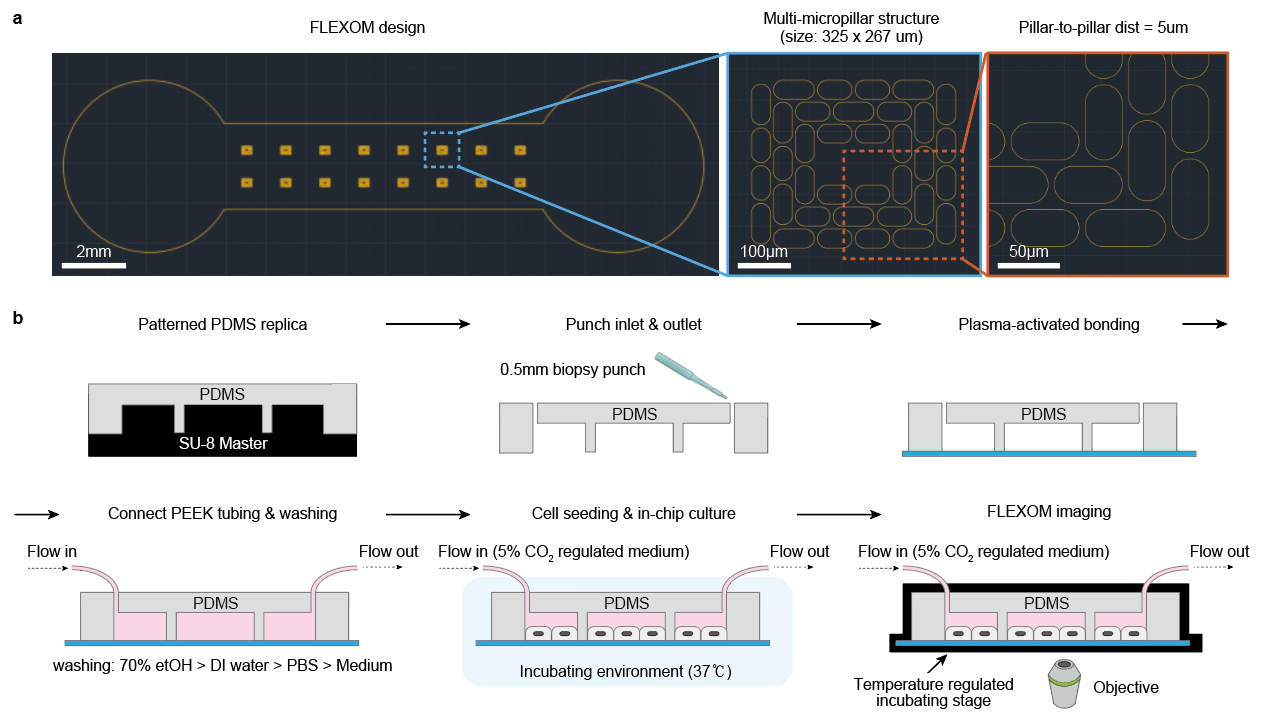

**Figure S1.** Design and experimental workflow of the FLEXOM microfluidic platform. a, CAD layout of the FLEXOM design, featuring inlet and outlet ports and a central culture chamber (left). Progressively magnified views detail the multi-micropillar structure (blue dashed box; middle) and the 5 µm pillar-to-pillar distance (orange dashed box; right). Scale bars, 2 mm (left), 100 µm (middle), and 50 µm (right). b, Schematic of the FLEXOM fabrication and experimental workflow, spanning PDMS replica molding, chip assembly, cell seeding, and live-cell imaging under continuous perfusion. See Methods for additional details.

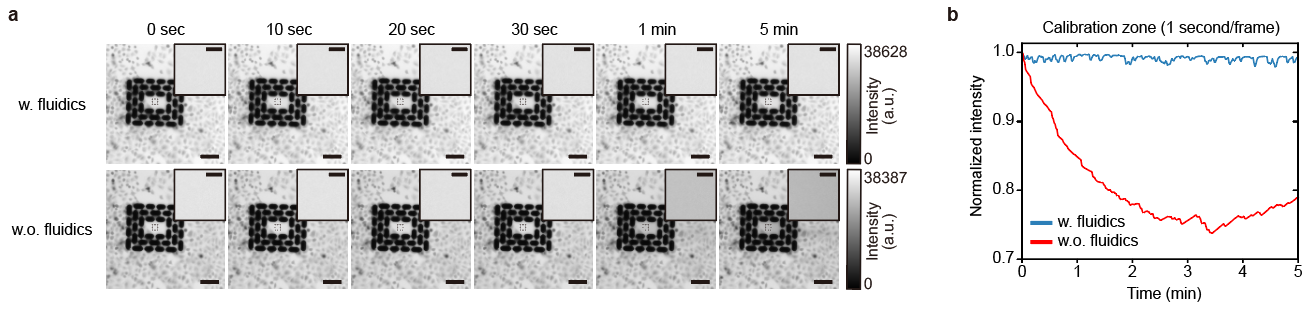

**Figure S2.** Fluidic exchange reduces photobleaching-driven fluorescence drift. **a,** Time-lapse fluorescence images acquired with (top) and without (bottom) continuous fluidic flow (*ΔP* = 1 kPa). Black dashed line, calibration zone; black box, magnified inset. Scale bars, 100 μm (main), 10 μm (inset). **b,** Mean fluorescence intensity of the calibration zone in a, normalized to the initial value, with (blue) and without (red) continuous fluidic exchange.

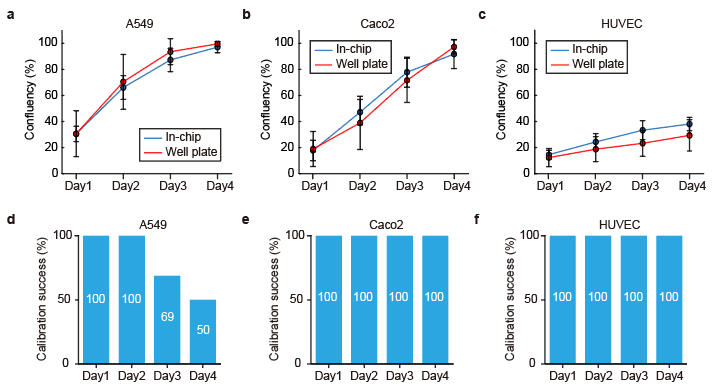

**Figure S3.** Biocompatibility and in situ self-calibration robustness of FLEXOM microfluidic chip. a–c, Confluency over 4 days for A549 (a), Caco-2 (b), and HUVEC (c) cells cultured in the FLEXOM chip (blue; n = 8 regions) or in standard 24-well plates (red; n = 36 regions); mean ± s.d. d–f, Percentage of regions with an available in situ calibration zone over 4 days for A549 (d), Caco-2 (e), and HUVEC (f); percentages are annotated in each bar (n = 16 regions). A calibration zone was classified as available when at least one cell-free 5 × 5 pixel zone for intensity referencing was present within the void region.

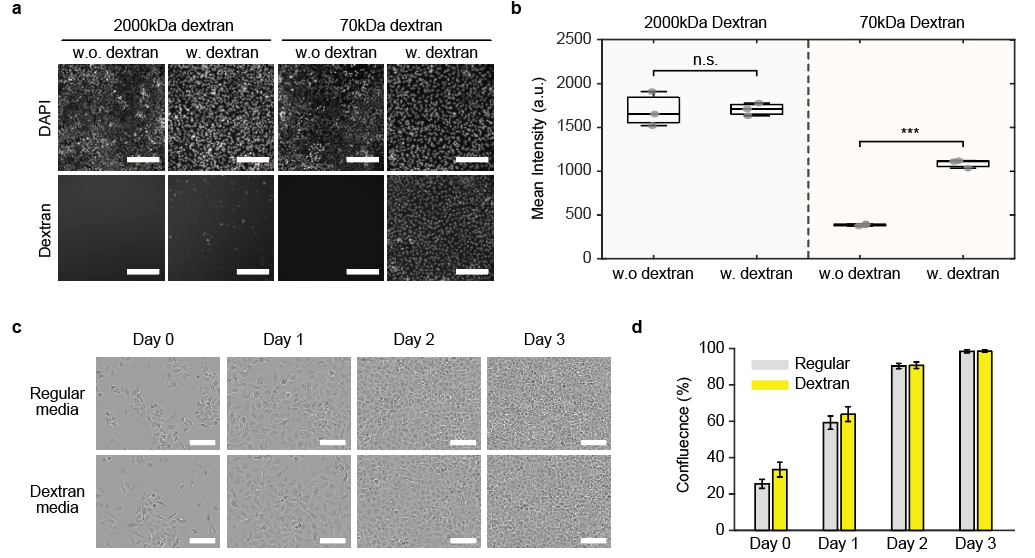

**Figure S4.** Cellular uptake and biocompatibility assessment of FITC-dextran. a, Fluorescence images of A549 cells incubated for 24 h with (w.) or without (w.o.) 2,000 kDa FITC-dextran or 70 kDa TRITC-dextran. Scale bars, 200 μm. b, Mean fluorescence intensity of dextran signal from a (n = 3 images per group). Statistical comparisons (two-sided Welch's t-test) were made between groups with and without dextran exposure. Box plots: median (line), IQR (box), min–max (whiskers); gray dots, individual data points. n.s., P > 0.05; ***P < 0.001. c, Phase-contrast images of cells exposed to 2,000 kDa FITC-dextran or regular medium over 3 days. Scale bars, 100 μm. d, Confluency over 3 days with (yellow) or without (gray) 2,000 kDa FITC-dextran (n = 36 regions of interest per group). Data depict mean (colored bars) ± s.d. (error bars).

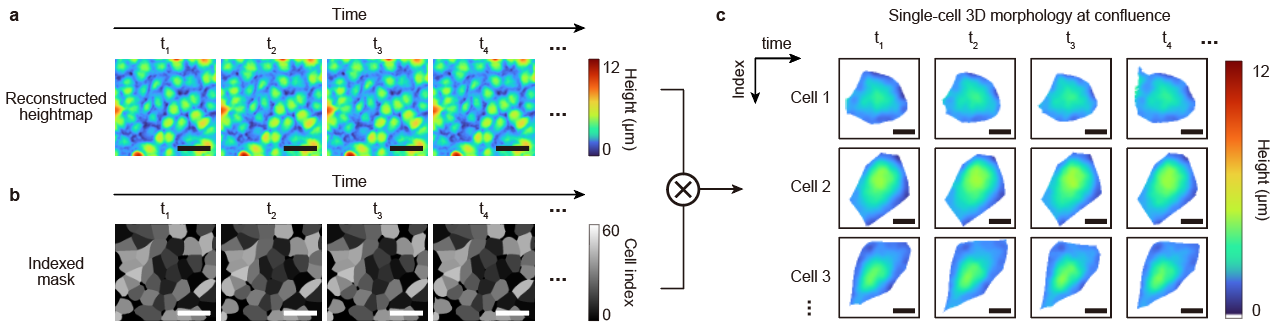

**Figure S5.** Single-cell 3D morphology reconstruction in a confluent monolayer. Pipeline for single-cell 3D reconstruction in confluent monolayers. a, Sequential FLEXOM height maps acquired over time. Scale bars, 50 μm. b, Cellpose-generated indexed cell masks with unique per-cell labels, propagated across time frames by TrackMate-based mask linkage to maintain consistent cell identities. Scale bars, 50 μm. c, Single-cell 3D morphology reconstructed by pixel-wise height assignment to labeled masks, yielding morphological parameters for each cell such as projected area, aspect ratio, and mean height (Note S1). Scale bars, 10 μm.

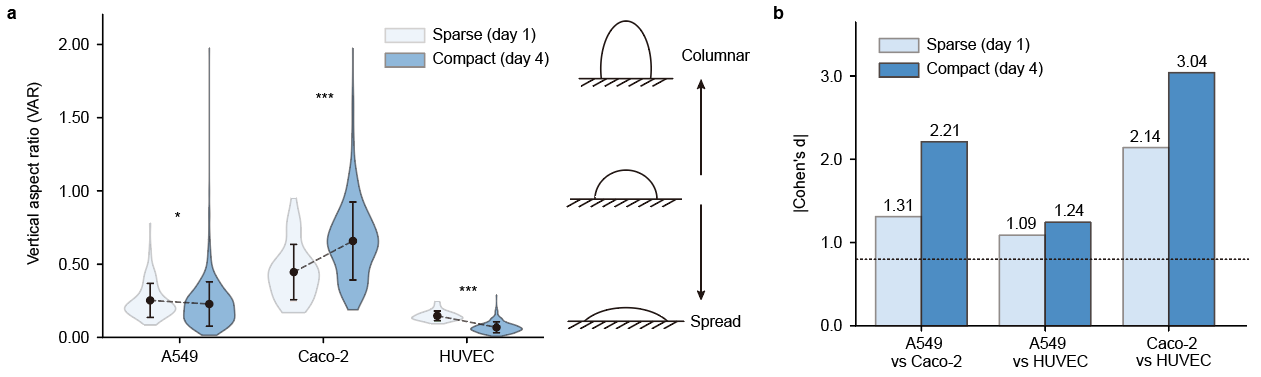

**Figure S6.** Cell-type-specific vertical morphology is amplified upon monolayer formation. a, Left, vertical aspect ratio (defined as mean height × area^-1/2^; VAR) of A549, Caco-2, and HUVEC cells in sparse (light blue; n = 124, 70, and 64 cells, respectively) and compact (dark blue; n = 885, 406, and 375 cells, respectively) conditions. Data depict mean (black dots) ± s.d. (error bars), with the slope of change (black dashed lines) from sparse to compact conditions. Statistical comparisons (two-sided Welch's t-test) between sparse and compact conditions yielded P = 3.5 × 10-2, 5.9 × 10-13, and 1.7 × 10-30 for A549, Caco-2, and HUVEC, respectively. *P < 0.05, ***P < 0.001. Right, schematic illustrating the relationship between VAR and vertical cell shape: high VAR corresponds to columnar morphology, low VAR to spread morphology. b, Effect sizes (|Cohen's d|; values above each bar) for pairwise cell-type comparisons of VAR in sparse (light blue) and compact (dark blue) conditions. The dashed line indicates the threshold for a large effect (|Cohen's d| = 0.8).

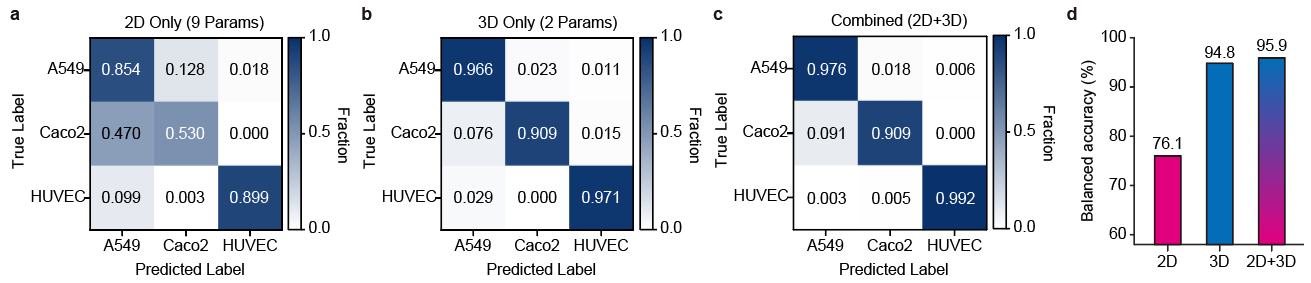

**Figure S7.** Classification performance using 2D and 3D morphological features. a–c, Confusion matrices for A549 (n=885), Caco-2 (n=406), and HUVEC (n=375) cells classified using 2D-only (a), 3D-only (b), and combined 2D+3D (c) feature sets, evaluated by 5-fold cross-validation. 2D features: area, aspect ratio, circularity, solidity, eccentricity, extent, perimeter, major axis, minor axis; 3D features: mean cell height and height standard deviation. d, Balanced classification accuracy per feature set (2D, 76.1%; 3D, 94.8%; 2D+3D, 95.9%). See Note S1 for additional details.

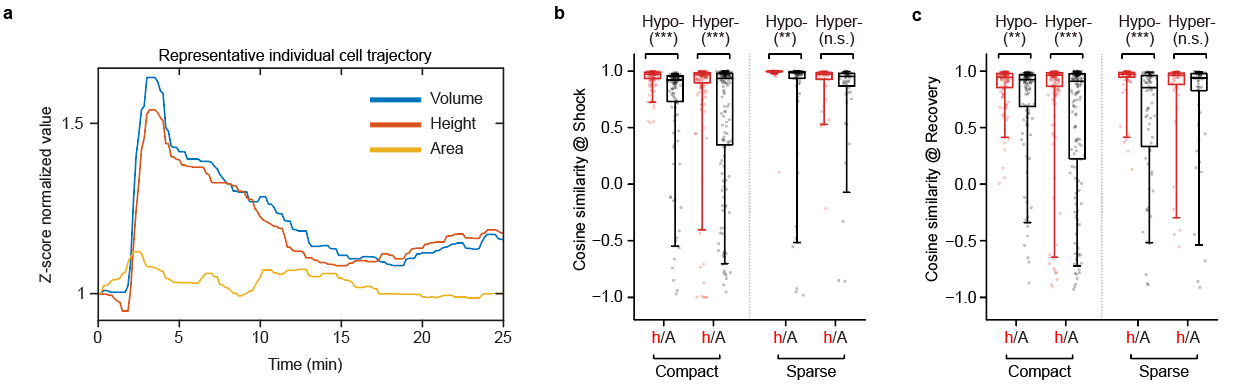

**Figure S8.** Correlation of morphological trajectories during osmotic shock. a, Representative Z-score normalized time-series trajectories of volume (blue), mean height (red), and projected area (yellow) for an individual cell. b,c, Cosine similarities of height (h; red) and area (A; black) trajectories relative to volume during the shock (b) and recovery (c) phases. Data are shown for compact monolayers under hypotonic (Hypo-; n = 120 cells; P = 9.8 × 10-7 and 2.8 × 10-3 for shock and recovery, respectively) and hypertonic (Hyper-; n = 165 cells; P = 9.2 × 10-5 and 2.6 × 10-5) conditions, and for sparse cells under hypotonic (n = 71 cells; P = 3.0 × 10-3 and 7.3 × 10-6) and hypertonic (n = 51 cells; P = 0.14 and 0.29) conditions. Box plots denote the interquartile range (box hinges) and 5th/95th percentiles (whiskers). Each dot represents one cell. Statistical comparisons between height- and area-based cosine similarities were performed using two-sided paired t-tests. **P < 0.01; ***P < 0.001; n.s., not significant.

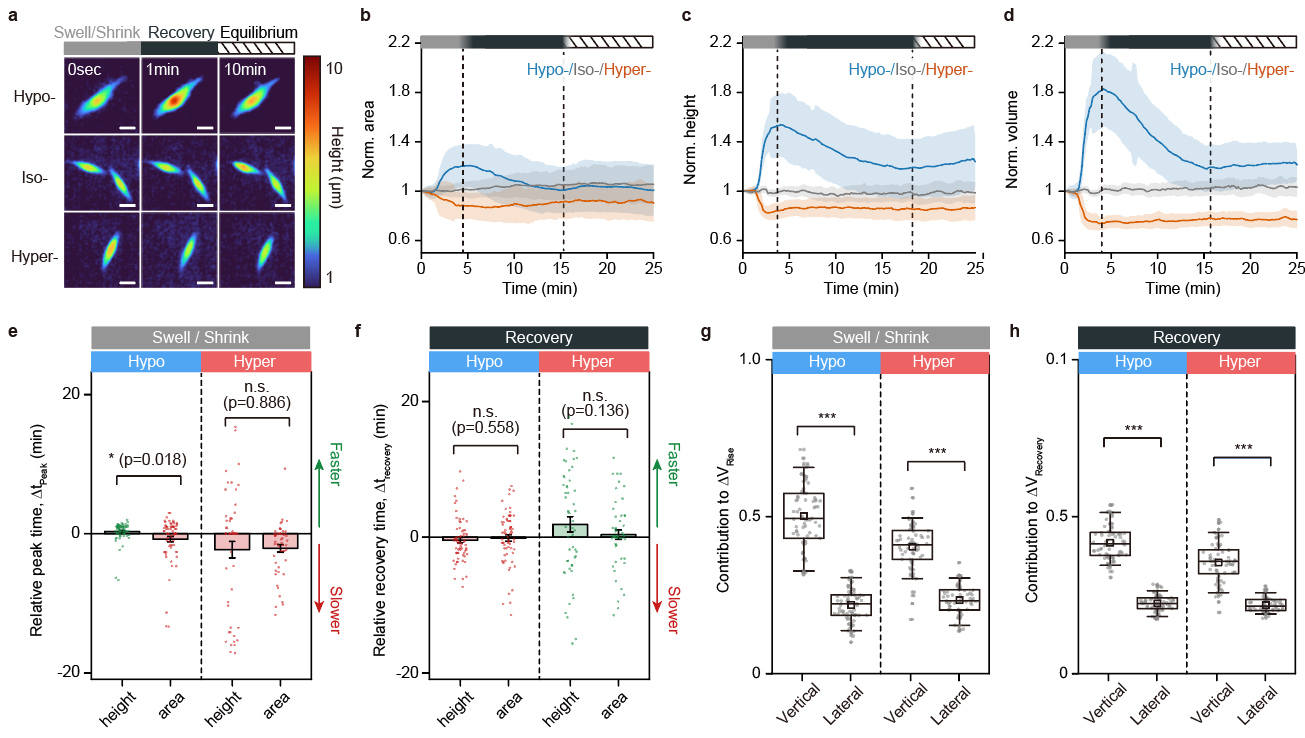

Figure S9. Osmotic responses in sparse (isolated) cells. a, Representative height maps of isolated A549 cells at 0 s, 1 min, and 10 min after osmotic perturbation under hypotonic (150 mOsm L-1; Hypo-), isotonic (330 mOsm L-1; Iso-), and hypertonic (450 mOsm L-1; Hyper-) conditions. Color bars indicate response stages: swelling/shrinking (gray), recovery (black), and equilibrium (striped). Scale bars, 20 μm. b–d, Normalized area (b), height (c), and volume (d) under Hypo- (blue), Iso- (gray), and Hyper- (red) conditions; mean ± s.d. Dashed vertical lines and color bars indicate transitions between response phases. Color-coding is defined as in a. e,f, Relative peak time (Δtpeak; e) and relative recovery time (Δtrecovery; f) for mean height and projected area during the swell/shrink and recovery phases (n = 67 and 52 cells for hypo- and hypertonic conditions, respectively). Data depict mean (colored boxes) ± s.d. (error bars). g,h, Vertical and lateral contributions to volume change up to peak (g) and throughout recovery (h); n = 71 and 65 cells for hypo- and hypertonic conditions, respectively. Box plots denote the interquartile range (box hinges), means (squares), 5th and 95th percentiles (whiskers). e-f, Statistical tests: two-sided paired t-test (within cells). n.s., not significant; *P < 0.05; ***P < 0.001.

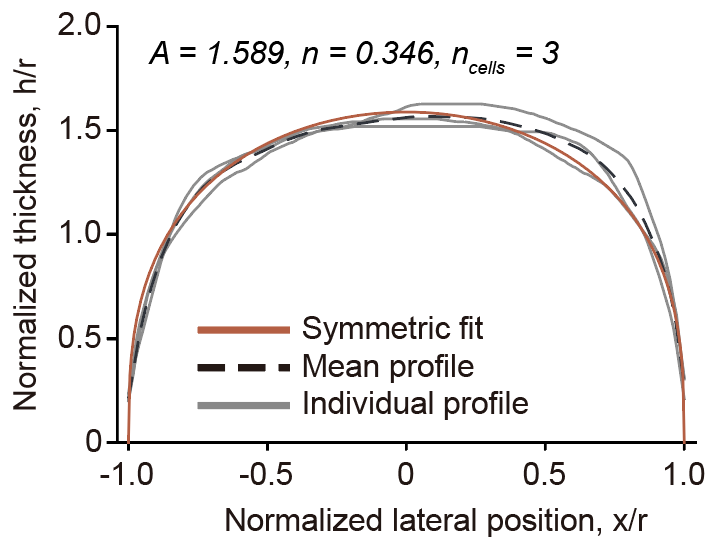

**Figure S10.** Normalized thickness profiles were extracted from individual cell masks (gray), interpolated onto a common grid, and averaged (black dashed). The mean profile was fitted to a symmetric dome function (brown; A = 1.589, n = 0.346; n = 3 masks).

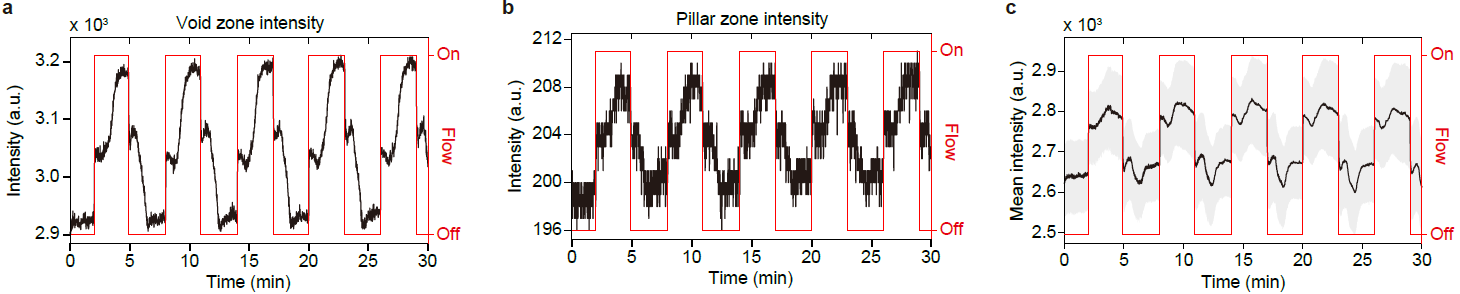

**Figure S11.** Fluorescence intensity of reference zones and individual cells during cyclic osmotic perturbations. a,b, Fluorescence intensity (black line) in the void zone (a) and pillar zone (b), used as frame-specific calibration references for height reconstruction during cyclic osmotic perturbations over 30 min (250 ms per frame). c, Mean fluorescence intensity of individual A549 cells in monolayers (n = 87 cells for N = 3 regions of interest) over the same imaging period. Right axes (red) indicate flow state (On, hypotonic; Off, isotonic). Data in c depict mean (black line) ± s.d. (gray shading).

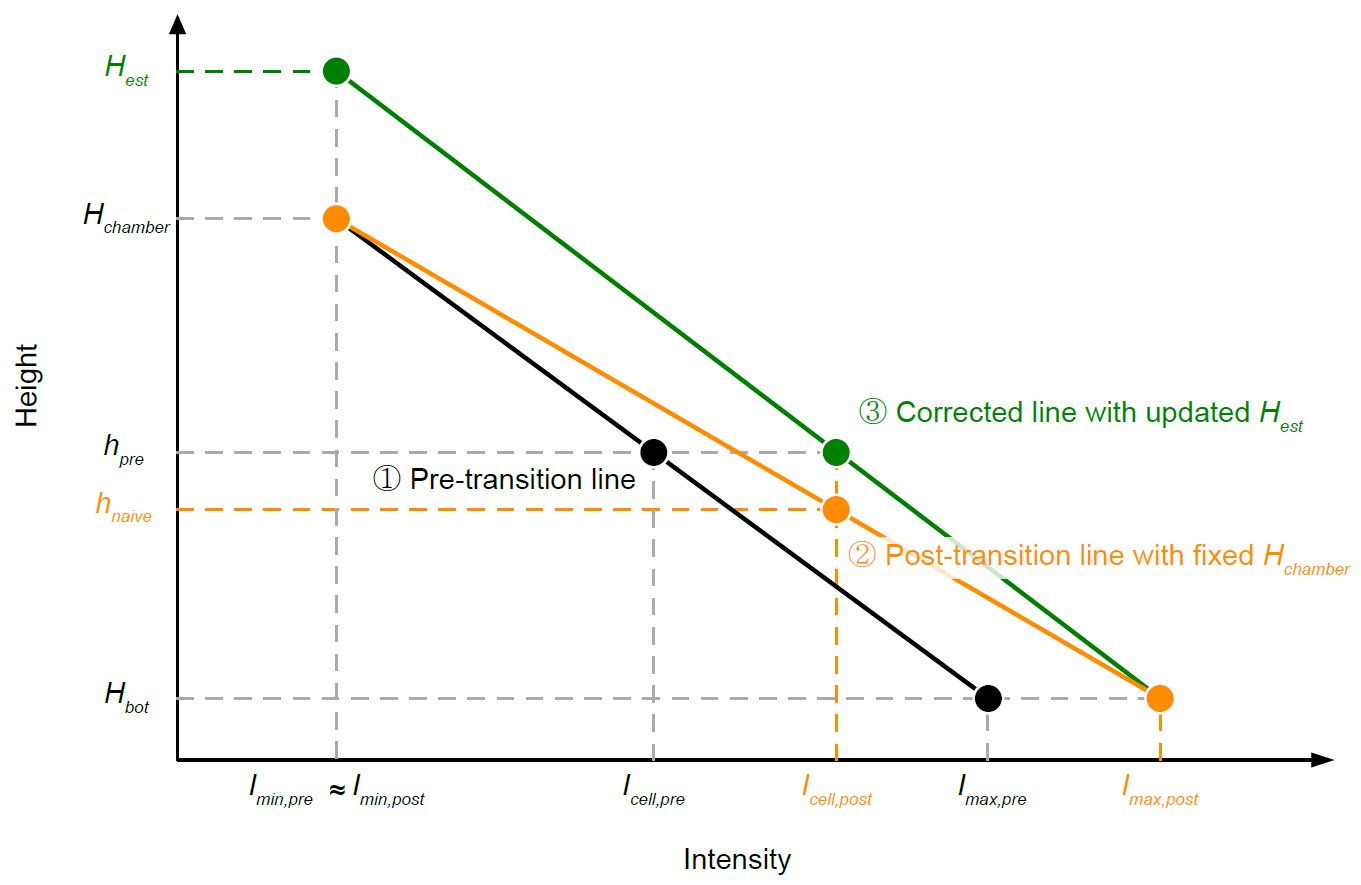

**Figure S12.** Schematic illustrating the three-step calibration correction applied after each stepwise intensity transition during cyclic osmotic perturbations. Pre-transition calibration line (black, ①) mapping cell fluorescence intensity (*I_cell,pre_*) to the current cell height (*h_pre_*) under the nominal chamber ceiling height (*H_chamber_*), anchored at the shared lower reference level (*H_bot_*). Post-transition line with fixed *H_chamber_* (orange, ②), which maps the post-transition cell intensity (*I_cell,post_*) to a lower apparent height (*h_naive_*) due to the shifted calibration endpoints (*I_max,post_* > *I_max,pre_*). Corrected post-transition line re-parameterized by the estimated ceiling height (*H_est_* ; green, ③), so that *I_cell,post_* maps consistently to *h_pre_*, with *H_bot_* and *I_min,pre_* ≈ *I_min,post_* unchanged across the transition.

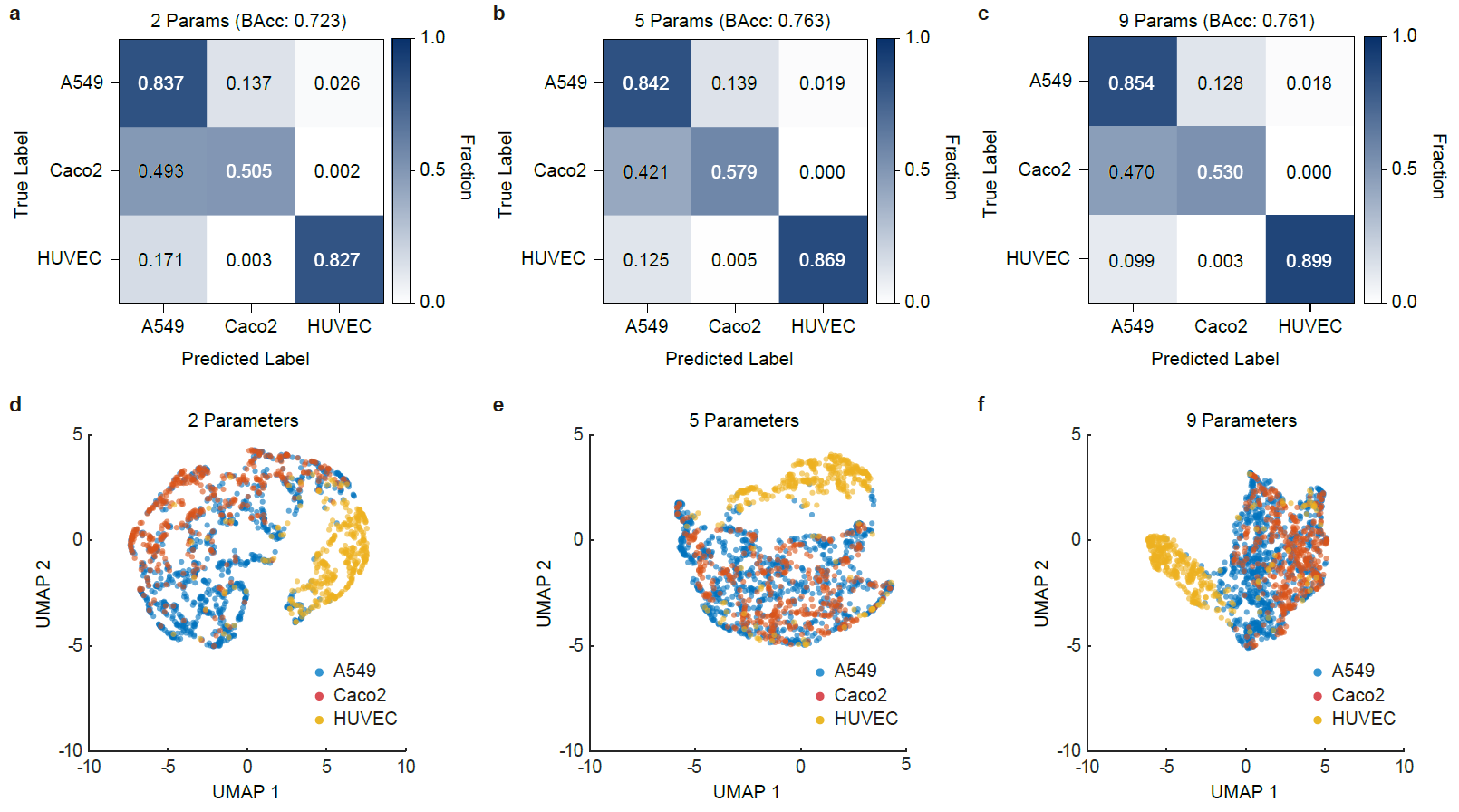

**Figure S13.** Cell-type classification performance using reduced 2D descriptor sets. a–c, Confusion matrices from 5-fold cross-validated linear SVM classifiers trained on 2D-only feature sets of 2 (a), 5 (b), and 9 (c) parameters (A549, n = 885; Caco-2, n = 406; HUVEC, n = 375). Balanced accuracies are 0.723, 0.763, and 0.761, respectively. d–f, Corresponding UMAP^[1]^ embeddings computed from the same 2D-only feature sets for visualization.

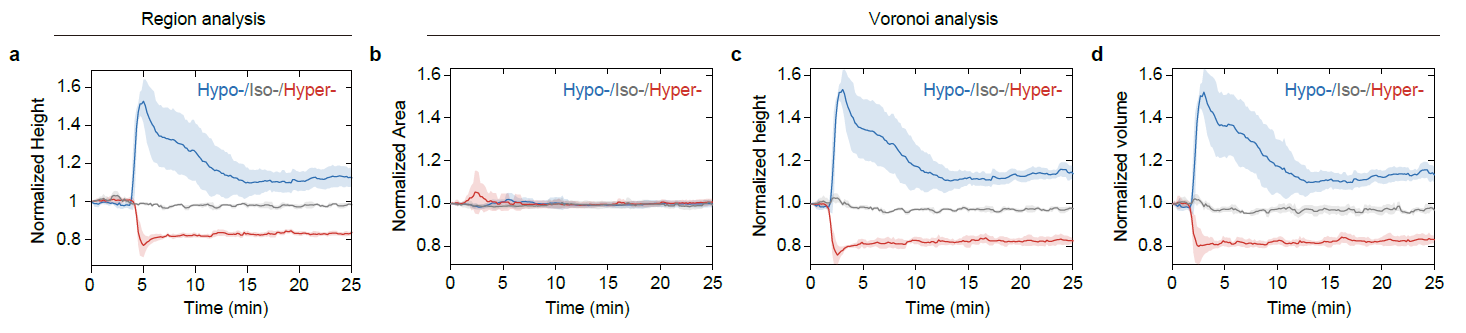

**Figure S14.** Consistency of osmotic response dynamics across analysis methods. a, Region-based analysis of normalized cell height over 25 min under hypotonic (150 mOsm L^-1^; Hypo-), isotonic (330 mOsm L^-1^; Iso-), and hypertonic (450 mOsm L^-1^; Hyper-) conditions (n = 7, 4, and 7 regions; N = 3, 2, and 3, respectively). b–d, Voronoi-based single-cell analysis of normalized projected area (b), height (c), and volume (d) under the same conditions, extracted from the identical field-of-view regions as in a (n = 103, 89, and 164 cells for Hypo-, Iso-, and Hyper- conditions, respectively). Data in a–d are mean ± s.d.

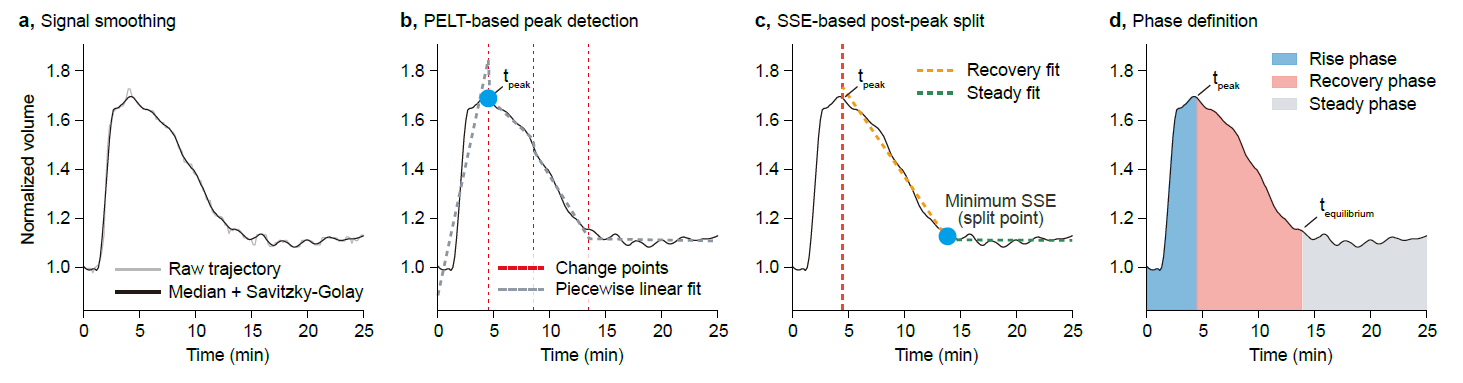

**Figure S15.** Automated pipeline for trajectory phase segmentation. a, Signal smoothing: raw trajectory (gray) and the output after 5-frame moving median followed by Savitzky–Golay filtering (black). b, Pruned Exact Linear Time (PELT)-based peak detection: the identified peak time (t_peak_; blue dot) is marked on the smoothed trajectory. Red dashed vertical lines indicate PELT-detected changepoints and the gray dashed line shows the resulting piecewise linear fit. c, Sum of squared errors (SSE)-based post-peak split: the two-segment linear fit minimising SSE separates the recovery (orange dashed) and steady-state (green dashed) phases; the optimal split point (blue dot) is indicated. d, Phase definition: rise phase (0 to t_peak_; blue), recovery phase (t_peak_ to t_equilibrium_; pink), and steady-state phase (t_equilibrium_ to end; gray).

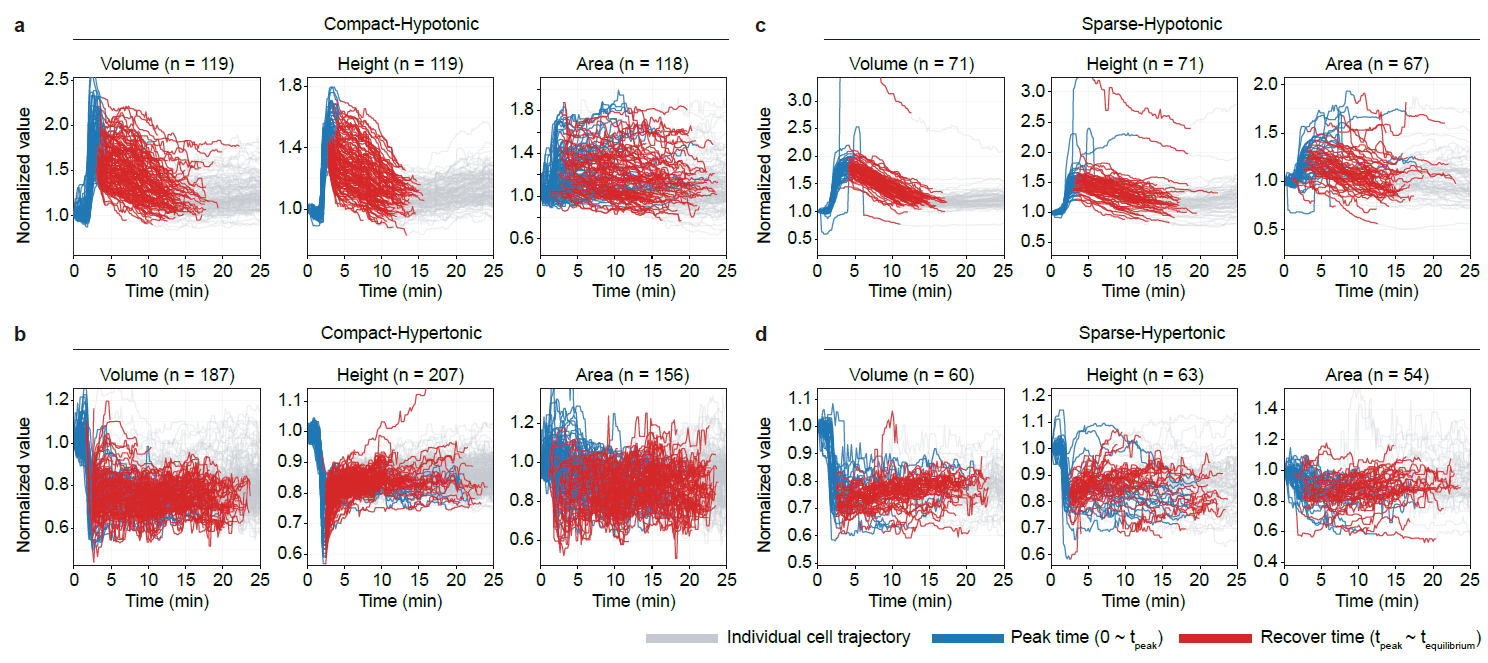

**Figure S16.** Individual cell trajectory segmentation during osmotic shock. Temporal profiles of normalized volume, height, and area for individual cells under four conditions, with values normalized to the initial time point (t = 0 min). a, compact monolayer, hypotonic; b, compact monolayer, hypertonic; c, sparse cells, hypotonic; d, sparse cells, hypertonic. Individual trajectories are shown in gray. Colored overlays indicate the rise phase (red; 0 to t_peak_) and recovery phase (blue; t_peak_ to t_equilibrium_), as identified by the pipeline in Figure S15. n is the number of cells with complete trajectories for each parameter.

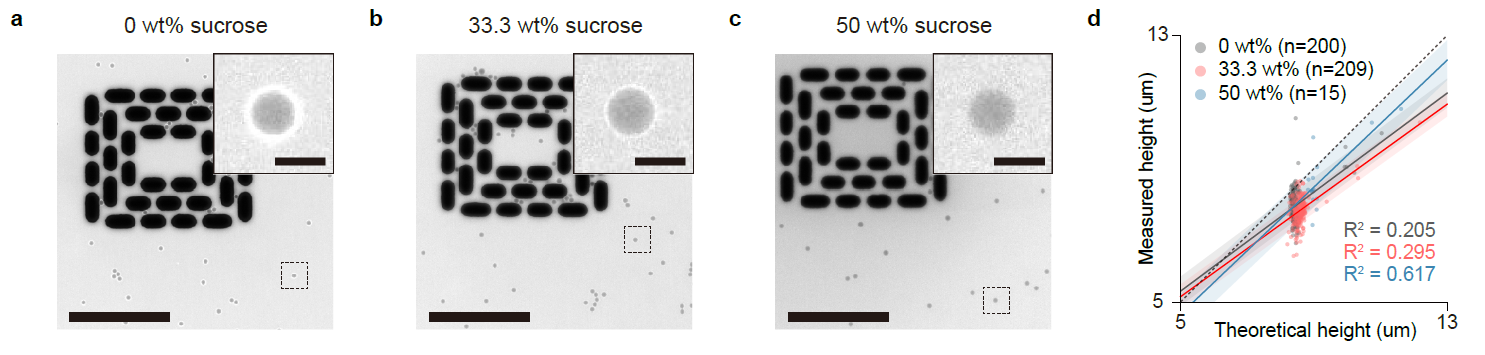

**Figure S17.** Fluorescence calibration accuracy as a function of refractive index (RI) matching. a–c, Representative wide-field fluorescence images of silica beads immersed in standard culture medium (0 wt% sucrose, RI ≈ 1.333; a), and in media supplemented with 33.3 wt% (RI ≈ 1.382; b) or 50 wt% sucrose (RI ≈ 1.398; c). Insets (dashed boxes in main images) show magnified views of individual beads. Scale bars, 200 μm (main), 50 μm (insets). d, Linear regression between theoretical and FLEXOM-measured bead heights at 0 wt% (gray), 33.3 wt% (red), and 50 wt% (blue) sucrose (n = 200, 209, and 15 beads, respectively).

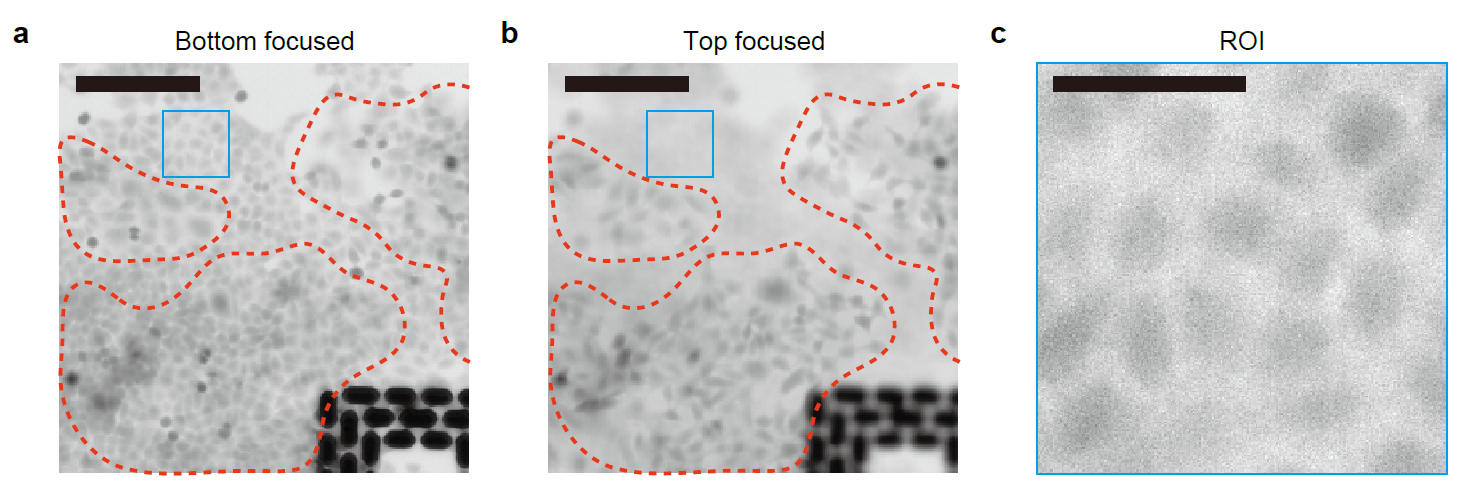

**Figure S18.** Exclusion of ceiling-attached cells prevents overestimation of cell thickness. a,b, Representative FLEXOM images focused on the bottom (a) and top (b) planes of the microfluidic chamber. Red dashed outlines indicate regions containing cells adhered to the chip ceiling, which were excluded from analysis. In these regions, signals from ceiling-attached and basal cells are superimposed within the same pixel column. Scale bars, 200 μm. c, Magnified view of the valid region of interest (ROI; blue box in a and b) selected from an area devoid of ceiling-attached cells. Scale bars, 50 μm.

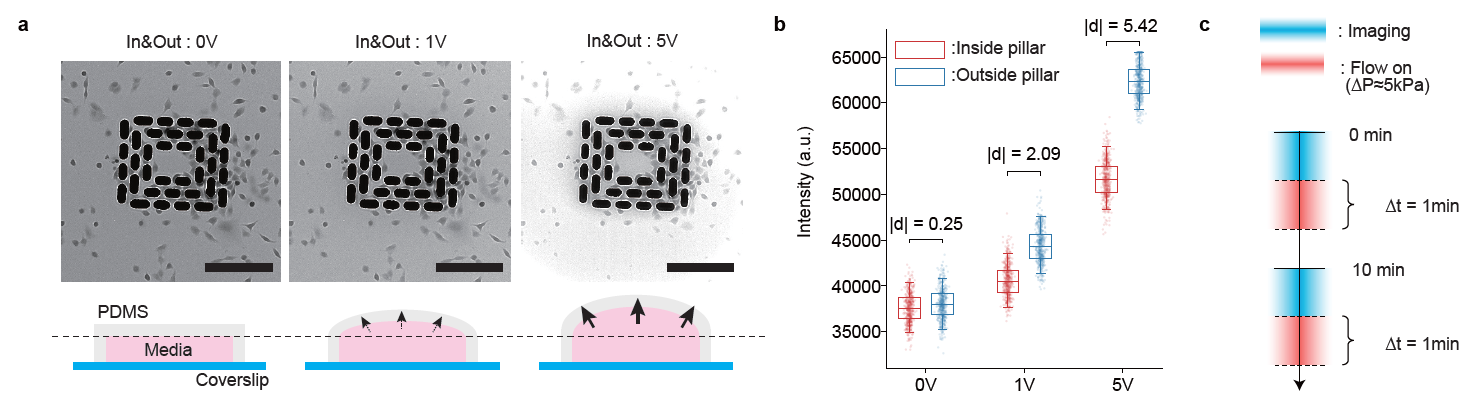

**Figure S19.** Pressure-induced chamber deformation and corresponding intensity offset. a, Representative fluorescence images (top) and corresponding cross-sectional schematics (bottom) of the microfluidic chamber under hydrostatic pressures of 0, 1, and 5 V applied simultaneously to the inlet and outlet. Black arrows indicate the direction of PDMS ceiling deflection. Scale bars, 200 μm. b, Fluorescence intensity of cell-free regions inside (red) and outside (blue) the multilayer micropillar structure at each pressure setting. Box plots denote the interquartile range (box hinges), median (line), and 5th and 95th percentiles (whiskers). Each dot represents one pixel measurement (n = 10,000 per group). Effect sizes (|Cohen's d|; values above brackets) quantify the difference between in- and outside the structure at each voltage. c, Experimental timeline for decoupled imaging and perfusion: differential pressure is maintained at 0 V during image acquisition (blue) and briefly elevated (ΔP ~ 5 kPa) for 1-min perfusion pulses between imaging intervals (red).

**Supplementary Note**

**Note S1 - Morphometric feature design and cell-type classification**

**Morphometric feature vector**

Cell boundaries were identified using Cellpose [2] and propagated across frames via the TrackMate [3] plugin in Fiji [4]. Each segmented cell $i$ is represented by an 11-dimensional morphometric feature vector

$$\mathbf{x}_{i}=\left[ x_{i,1},\ldots,x_{i,11} \right]\in\mathbb{R}^{11}$$

comprising nine 2D lateral descriptors (indices $1$–$9$) and two 3D height descriptors (indices $10$–$11$). The 2D descriptors quantify the geometry of the binary footprint mask; the 3D descriptors summarize the apical height profile within that region.

The 2D descriptors each capture one aspect of size, elongation, or shape compactness:

$$\begin{matrix} x_{i,1} & =\mathrm{AR}_{i}=\frac{L_{i}^{\mathrm{maj}}}{L_{i}^{\min}} \text{(aspect ratio)}, \\ x_{i,2} & =A_{i} \text{(projected area)}, \\ x_{i,3} & =\mathrm{Circ}_{i}=\frac{4\pi A_{i}}{P_{i}^{2}} \text{(circularity)}, \\ x_{i,4} & =\mathrm{Sol}_{i}=\frac{A_{i}}{A_{i}^{\mathrm{convex}}} \text{(solidity)}, \\ x_{i,5} & =\mathrm{Ecc}_{i} \text{(eccentricity)}, x_{i,6}=\mathrm{Ext}_{i} \text{(extent)}, \\ x_{i,7} & =P_{i} \text{(perimeter)}, x_{i,8}=L_{i}^{\mathrm{maj}} \text{(major axis)}, x_{i,9}=L_{i}^{\min} \text{(minor axis)}, \end{matrix}$$

where $A_{i}$ is the projected area, $P_{i}$ is the perimeter, and $L_{i}^{\mathrm{maj}}$, $L_{i}^{\min}$ are the major and minor axis lengths of the best-fit ellipse. Denoting the reconstructed height map as $h\left( x,y \right)$ and the projected domain of cell $i$ as $\Omega_{i}$, the 3D descriptors are:

$$\begin{matrix} x_{i,10} & =\bar{h}_{i}=\langle h\rangle_{\Omega_{i}} \text{(mean height)}, \\ x_{i,11} & =\sigma_{h,i}=std\left( h \right)_{\Omega_{i}} \text{(height s.d.)}. \end{matrix}$$

Cell volume, used separately for osmotic response analysis (Note S2), was not included in the classification feature vector because it is a derived quantity ($V_{i}\approx A_{i}\cdot\bar{h}_{i}$) that combines lateral and vertical information already captured by the individual descriptors. Because the feature vector spans heterogeneous units, each dimension was $z$-scored using statistics computed from the training folds only to prevent data leakage.

**Classification and accuracy metrics**

Cell-type prediction was performed using a linear support vector machine (SVM) [5] with $\mathcal{l}_{2}$ regularization and an error-correcting output codes (ECOC) [6] strategy, evaluated by 5-fold stratified cross-validation. Balanced accuracy (BAcc) [7]—the macro-average of per-class recall—was used as the primary metric to account for class-size imbalance (Figure S7):

$$BAcc=\frac{1}{K}\sum_{u=1}^{K} \frac{C_{u,u}}{\sum_{v} C_{u,v}},$$

where $\mathbf{C}$ is the $K\times K$ confusion matrix, with $C_{u,v}$ denoting the number of cells of true class $u$ predicted as class $v$.

**Feature ablation: 2D vs. 3D morphology**

To determine which morphology channel carries discriminative information, classifiers were trained on the following feature subsets (Figure S7, Figure S13):

$$\begin{matrix} {\tilde{\mathbf{x}}}_{i}^{\left( 2D \right)} & =\left[ \tilde{x}_{i,1},\ldots,\tilde{x}_{i,9} \right]\in\mathbb{R}^{9} \text{(BAcc}=0.761\text{)}, \\ {\tilde{\mathbf{x}}}_{i}^{\left( 3D \right)} & =\left[ \tilde{x}_{i,10},\tilde{x}_{i,11} \right]\in\mathbb{R}^{2} \text{(BAcc}=0.948\text{)}, \\ {\tilde{\mathbf{x}}}_{i}^{\left( \mathrm{All} \right)} & =\left[ \tilde{x}_{i,1},\ldots,\tilde{x}_{i,11} \right]\in\mathbb{R}^{11} \text{(BAcc}=0.959\text{)}. \end{matrix}$$

Sensitivity analyses with reduced 2D subsets further showed that accuracy improved only incrementally with additional 2D descriptors (2 params [aspect ratio, area]: BAcc $=0.723$; 5 params [aspect ratio, area, circularity, solidity, eccentricity]: $0.763$; 9 params [all nine 2D descriptors]: $0.761$; Figure S13), confirming that increasing 2D descriptor count alone is insufficient for robust discrimination. By contrast, just two height-derived parameters ($\bar{h}$, $\sigma_{h}$) achieved BAcc $=0.948$, substantially outperforming all 2D configurations. These results demonstrate that vertical height encodes cell-type-specific morphological information largely orthogonal to 2D lateral geometry (Figure 3h,i).

**Representative 3D morphology reconstruction**

To visualize the characteristic geometry of each cell type, a representative synthetic surface was generated from group-averaged morphometric parameters (Figure 3f). The lateral boundary was modeled as an ellipse with semi-axes derived from the mean projected area $\langle A\rangle_{g}$ and mean aspect ratio $\langle AR\rangle_{g}$:

$$a_{g}=\sqrt{\frac{\langle A\rangle_{g} \langle AR\rangle_{g}}{\pi}}, b_{g}=\sqrt{\frac{\langle A\rangle_{g}}{\pi\langle AR\rangle_{g}}}.$$

The apical surface was parameterized as a dome using a normalized radial coordinate $\rho\in\left[ 0,1 \right]$:

$$H_{g}\left( \rho\right)=h_{0,g} \left( 1-\rho^{2} \right)^{q_{g}},$$

where the peak height $h_{0,g}=\langle\bar{h}\rangle_{g} \left( q_{g}+1 \right)$ ensures that the model mean height matches the empirical group mean. The shape exponent $q_{g}$, which controls apex sharpness, was determined by least-squares fitting to the observed area fraction $F_{obs,g}$, defined as the fraction of the cell footprint where height exceeds the group mean height. For the dome model, this fraction depends only on $q$:

$$F_{\mathrm{model}}\left( q \right)=1-\left( \frac{1}{q+1} \right)^{1/q},$$

and $q_{g}$ was chosen to minimize $\left| F_{\mathrm{model}}\left( q_{g} \right)-F_{obs,g} \right|$. The surface was mapped to Cartesian coordinates as $x=a_{g}\rho\cos\theta$, $y=b_{g}\rho\sin\theta$, $z=H_{g}\left( \rho\right)$, with axis limits held constant across all groups for fair visual comparison.

For phase-resolved visualization of osmotic responses (Figure 4c,d), the same parametric dome model was applied at each time point using per-minute group-averaged statistics ($\langle A\rangle_{g,t}$, $\langle AR\rangle_{g,t}$, $\langle\bar{h}\rangle_{g,t}$, and $F_{obs,g,t}$), generating a sequence of representative surfaces that capture the dynamic morphological response to osmotic perturbation.

**Note S2 - Anisotropic and height-dominated volume regulation during osmotic shock**

**Vertical bias of osmotic deformation**

Cross-sectional height profiles revealed that osmotic deformation was predominantly vertical (Figure 4c,d). In confluent monolayers, hypotonic shock increased mean cell height by 45.7% at peak (n = 120 cells), whereas hypertonic shock decreased mean cell height by 24.0% (n = 210 cells). Lateral area changed on a slower timescale (Figure 4e–h). Recovery was incomplete under both conditions, with residual height offsets of +13.9% after hypotonic shock and −14.3% after hypertonic shock relative to pre-shock baselines. These trends were consistent across region-based and single-cell Voronoi analyses (Figure S14), confirming robustness to the segmentation approach.

**Height as a temporal proxy: the influence of physical constraints**

We quantified the divergence in response kinetics between mean cell height (h̄) and projected area (A) using an automated trajectory segmentation pipeline (Figure S15). First, raw trajectories were smoothed with a moving median filter (window size = 5 time points) to suppress transient outliers, followed by a second-order Savitzky–Golay filter [9] (window size = 11 time points, mode = interpolation) to preserve the overall trajectory shape (Figure S15a). Next, a two-dimensional signal comprising the smoothed values and corresponding time points was submitted to the Pruned Exact Linear Time (PELT) change-point detection algorithm [10] (ruptures, linear cost model; minimum segment size = 8, jump = 2). A BIC-inspired base penalty was computed as σ̂² ln(n), where σ̂² is the variance of the smoothed trajectory and n is the number of time points. Because no single penalty universally optimized segmentation across cells, the algorithm was run at five penalty multipliers (8, 12, 16, 20, and 24 times the base penalty), and the segmentation that best satisfied a composite criterion, valid peak position, segment count closest to four, and lowest mean squared error, was retained. The peak response time (t_peak_) was then defined as the time of the piecewise-linear maximum for hypotonic stimulation or minimum for hypertonic stimulation in the selected segmentation (Figure S15b). The post-peak trajectory was split at the time point that minimized the summed squared error (SSE) of a two-segment linear fit (minimum segment length = 5 time points), partitioning the response into relaxation and steady-state phases (Figure S15c,d). Individual cell trajectories segmented into these three phases are shown in Figure S16. For paired temporal comparisons, only cells for which valid rise and recovery metrics were obtained for all three variables, volume, height, and projected area, were retained as common cells (confluent: n = 116 hypotonic cells and 152 hypertonic cells; isolated: n = 67 hypotonic cells and 52 hypertonic cells).

Height consistently peaked and began recovering earlier than projected area in confluent monolayers (Figure 4k,l; two-sided paired t-test, P < 0.001). Within a confluent monolayer, cells are physically constrained by their neighbors: while height can adjust relatively freely into the aqueous medium above, lateral changes require the coordinated displacement or deformation of the surrounding cellular network [11]. We attribute the observed temporal lag in lateral remodeling to this mechanical coupling and frictional resistance at cell–cell junctions [11].

**Height as the primary driver: mechanical efficiency of vertical remodeling**

To determine whether height is the dominant physical contributor to volume change, we decomposed volume dynamics along the lateral and vertical axes. Defining an effective lateral length $X_{i}\left( t \right)=\sqrt{A_{i}\left( t \right)}$ yields the logarithmic identity:

$$\Delta\ln V_{i}\left( t \right)=2 \Delta\ln X_{i}\left( t \right)+\Delta\ln\bar{h}_{i}\left( t \right),$$

where $\Delta$ denotes the difference between successive time steps. To quantify the relative contribution of each axis, instantaneous fractional contributions were defined as:

$$C_{vert,i}\left( t \right)=\frac{\left| \Delta\ln\bar{h}_{i}\left( t \right) \right|}{\left| \Delta\ln\bar{h}_{i}\left( t \right) \right|+2 \left| \Delta\ln X_{i}\left( t \right) \right|+\epsilon}, C_{lat,i}\left( t \right)=\frac{\left| \Delta\ln X_{i}\left( t \right) \right|}{\left| \Delta\ln\bar{h}_{i}\left( t \right) \right|+2 \left| \Delta\ln X_{i}\left( t \right) \right|+\epsilon},$$

where ε is machine epsilon to avoid division by zero. The observed C_vertical_ consistently exceeded C_lateral_ across both phases and both conditions (confluent hypotonic: 0.459 ± 0.010 vs. 0.220 ± 0.005 at peak, 0.419 ± 0.005 vs. 0.219 ± 0.002 during recovery; confluent hypertonic: 0.450 ± 0.006 vs. 0.229 ± 0.003 at peak, 0.349 ± 0.003 vs. 0.230 ± 0.002 during recovery; two-sided paired t-test, P < 0.001 for all comparisons; Figure 4i,j).

**Trajectory correlation analysis**

To independently assess which morphological parameter more faithfully tracks volume dynamics, we computed the cosine similarity between $z$-score-normalized time-series trajectories of volume and each morphological parameter for every cell. For cell $i$, the similarity between volume and height was defined as

$$S_{h,i} = \frac{\mathbf{v}_{i}\cdot\mathbf{h}_{i}}{\parallel\mathbf{v}_{i}\parallel\parallel\mathbf{h}_{i}\parallel},$$

where $\mathbf{v}_{i}$ and $\mathbf{h}_{i}$ denote the $z$-score-normalized volume and mean height trajectories, respectively; $S_{A,i}$ was defined analogously for the projected area trajectory. Each trajectory was evaluated over two non-overlapping temporal windows: the shock phase ($0$ to $t_{\mathrm{peak}}$) and the recovery phase ($t_{\mathrm{peak}}$ to $t_{\mathrm{equilibrium}}$). Under both hypotonic and hypertonic perturbations, height trajectories exhibited consistently higher cosine similarity to volume than did area trajectories (Figure S8).

**Comparison with isolated cells: the role of confluency**

To test whether height-dominated regulation arises from cell-intrinsic properties or from lateral mechanical confinement, we repeated the above analyses in isolated (sparse) cells. Consistent with prior work demonstrating that confluence and tight junction integrity are required for epithelial volume regulation [12], the temporal decoupling between height and area responses was substantially reduced: in sparse hypotonic cells, the difference in rise times between height and area was markedly smaller than under confluent conditions ($P=0.018$ vs. $P<{10}^{-15}$), and in sparse hypertonic cells, rise times were indistinguishable ($P=0.886$ vs. $P<{10}^{-15}$; Figure S9e,f; Figure S16c,d). However, the per-axis contribution analysis revealed that $C_{\mathrm{vert}}$ remained above the isotropic baseline of $1/3$ even in isolated cells (sparse hypotonic: $C_{\mathrm{vert}}=0.495$ at peak, $0.411$ during recovery; sparse hypertonic: $0.399$ at peak, $0.349$ during recovery; Figure S9g,h), indicating that a degree of vertical bias persists independently of junctional confinement.

**Note S3 - Measurement limitations and device constraints**

**Nonlinearity of height measurement at low cell heights**

Two non-exclusive mechanisms may account for the systematic deviation observed in Supplementary Note 1 at low cell heights ($\lesssim5 \mu m$). First, confocal-based height reconstruction requires segmentation of the actin-labeled boundary, which introduces uncertainty when the boundary is diffuse; the discrepancy should therefore not be attributed solely to FLEXOM error. Second, the intensity–height relation may depart from linearity due to the finite numerical aperture (NA) of the objective [13]. Under the ideal paraxial approximation, all collected rays travel vertically through the dye layer, so intensity scales linearly with dye-layer thickness, $I\left( z \right)\approx\alpha z$. In practice, a finite-NA objective collects fluorescence over an angular cone extending to $\theta_{\max}=arcsin\left( NA/n \right)$, so the total detected signal becomes

$$I\left( z \right)\propto z\int_{0}^{\theta_{\max}} W\left( \theta,z \right) \tan\theta d\theta,$$

where $W\left( \theta,z \right)$ is an effective angular weight. In the small-gap regime ($z\lesssim$ a few micrometers), optical confinement effects cause $W$ to acquire a $z$-dependence, causing the measured intensity to deviate from simple linear scaling.

**Refractive index mismatch and calibration nonlinearity**

An additional contributor to nonlinearity is refractive index (RI) mismatch between the imaging medium and the sample boundaries (glass, PDMS, and cell membrane) [13,14]. To quantify this effect, silica beads (RI $\approx$ 1.46) of known geometry were used as reference objects, with sucrose added at 0, 33.3, or 50 wt% to progressively raise the medium RI. The correlation between the theoretical and FLEXOM-measured bead heights improved with increasing RI matching ($R^{2}=0.205$ at 0 wt%; $R^{2}=0.295$ at 33.3 wt%; $R^{2}=0.617$ at 50 wt%; Figure S17d). All biological experiments were conducted in standard isotonic medium (RI $\approx$ 1.333), accepting the associated calibration nonlinearity as an inherent constraint of the current implementation.

**Inherent limitations of fluorescence exclusion-based thickness reporting**

Three measurement limitations arise from the fluorescence exclusion principle. First, FLEXOM reports the total vertical thickness of the dye-excluded region rather than the absolute $z$-position of a specific cellular surface. For suspended or mitotic cells that detach from the substrate, the reported thickness overestimates the true apical height. Second, vertical overlap between cells causes FLEXOM to report the combined exclusion thickness of all superimposed cell bodies within a single pixel column. Regions where cells adhered to the ceiling were therefore identified from top-focused images and excluded from all subsequent analyses (Figure S18). Third, FLEXOM cannot determine the absolute axial location of the junctional complex, though this limitation does not affect the height or volume measurements reported in this study.

**Device-level constraints**

Two device-specific factors impose practical limits on FLEXOM measurements. First, the microfluidic chamber height of $27 \mu m$ sets an upper bound on the measurable cell height. Second, the compliance of the PDMS ceiling introduces a flow-dependent artifact: application of hydrostatic pressure deflects the ceiling upward, increasing the dye volume and artifactually elevating fluorescence intensity (Figure S19a) [8]. This deflection is disproportionately large in regions outside the micropillar array, with effect sizes ($|$Cohen’s $d|$) between inside and outside the array increasing from $0.25$ at 0 V to $5.42$ at 5 V (Figure S19b). Two mitigation strategies are: (i) temporal decoupling of flow and imaging (Figure S19c); or (ii) pre-characterization of the pressure-dependent deformation profile by 3D confocal imaging.
